# Molecular dynamics simulations demonstrate reduced antibiotic affinity to mirror bacterial targets

**DOI:** 10.64898/2026.07.14.738450

**Authors:** Paul-Enguerrand Fady, Juliette Ciccone

**Affiliations:** The Centre for Long-Term Resilience, London WC2H 9JQ, United Kingdom; UCL Chemistry, Christopher Ingold Building, 20 Gordon Street London, WC1H 0AJ, United Kingdom

## Abstract

“Mirror life”, self-replicating organisms composed of non-natural-chirality biomacromolecules, presents a future threat with potentially global consequences. Consequently, there is strong agreement among experts that it should not be created. However, there is some disagreement over how effective existing medical countermeasures might prove against mirror bacteria in the event that they were created. Here, we leverage computational chemistry methods including docking and molecular dynamics to determine the likely binding efficacy of existing antibiotics against natural and mirror bacterial protein targets. We find that most existing antibiotics fail to bind to mirror bacterial protein targets, unlike their natural-chirality targets. This suggests altered binding of current medical countermeasures, which may impact the antimicrobial activity against mirror bacteria were the latter were created.

## 1. Introduction

Living organisms are composed of biomacromolecules with a specific chirality. All known life forms exhibit homochirality: glycans are predominantly composed of sugars in the D-configuration (such as D-glucose, D-mannose, D-galactose), i.e. sugars with right-handed chiral centres.^1^ The D 2 deoxyribose of DNA is also chiral, imparting a right-handed directionality to the helical twist.^2^ Conversely, proteins in living organisms are composed predominantly of L-amino acids, i.e. ones with left-handed chiral centres.^3^ While it is possible for monomers of the opposite chirality to be integrated into biomacromolecules (and even common in e.g. peptidoglycan bacterial cell walls which contain D-alanine and D-glutamate),^4^ the predominant pattern of chirality is fixed in one specific form for individual biomacromolecules.

“Mirror life”, or synthetic living organisms composed of biomacromolecules built using the non-naturally-occurring enantiomers of biomolecular monomers, presents an alternative option to this universal truth. Recent advances in synthetic cell biology have transformed mirror life from a theoretical science-fiction scenario to a material future threat. Synthesising life from matter remains an unsolved scientific problem which, if solved, would enable the creation of natural-chirality life as well as mirror life. While the exact timeline for the development of self-replicating mirror organisms is disputed, experts estimate that it could be achieved within the next 10 to 30 years, depending on resource allocation.^5^

Mirror life could represent a future threat with potentially global consequences. A technical report published in December 2024 outlines the ways in which mirror bacteria could subvert human, animal, and plant immune systems, leading to uncontrolled growth due to lack of recognition by stereospecific defences.^5^ Were mirror bacteria to be created and find their way into the environment – whether through deliberate release or a containment breach – they could proliferate extensively. This could ultimately lead to nutrition depletion in key ecological niches, starving keystone species of nutrients, which could drive severe downstream effects.

The environmental picture motivates policy recommendations to ban research that directly leads to the creation of mirror life. This is distinct from research which may involve mirror macromolecules, which must be permitted in order to foster innovation in the life and material sciences. There is widespread agreement among scientific experts from a range of domains that mirror life should not be created, as noted in the Policy Forum piece accompanying the 2024 technical report;^5^ the “Spirit of Asilomar” conference statement on mirror life in early 2025;^6^ the Paris Conference on Risks from Mirror Life Meeting Report from mid-2025;^7^ a commentary by the German Central Commission for Biological Safety;^8^ and a report from the UNESCO International Bioethics Committee.^9^

However, there has been pushback from some experts surrounding the actual level of risk associated with self-replicating mirror organisms, such as mirror bacteria. A small minority of dissenting voices have indicated their belief that current medical countermeasures such as antibiotics would successfully neutralise the threat of mirror organisms by virtue of their non-stereospecific activity.

There is early evidence that antibiotics would not be effective against mirror bacteria. In the absence of mirror bacteria against which to directly test compounds, two parallel strategies have emerged to determine the likely efficacy of current countermeasures. The first is to synthesise mirror-chirality antibiotics and test them against native bacteria. This strategy relies on the assumption of equivalency between the interaction of mirror antibiotics with native-chirality bacteria and native-chirality antibiotics with mirror bacteria.^10^ While this provides useful experimental evidence, it leverages costly enantiomers of antibiotics and requires “wet lab” resources which constrain the throughput of experiments.

The other strategy is to investigate antibiotic efficacy against mirror bacteria utilising computational chemistry. Thus far, docking and molecular dynamics have been used to investigate the efficacy of amoxicillin against native-chirality and mirror bacterial protein targets from *Chlamydia trachomatis* and *Staphylococcus aureus*.^11,12^ Computational approaches enable investigations into this sensitive topic in a risk-free manner, i.e. without generating any mirror components which may contribute to development of mirror life. In addition, this strategy provides a reproducible and iterative workflow which can be cheaply and rapidly adapted to study a range of targets with minimal resource requirements. This provides a risk-free environment for high-throughput determination of binding efficacy. A reduced binding efficacy may plausibly have a negative effect on antimicrobial activity – though this work is strictly computational and does not present *in vitro* validation of results.

In this work, we have chosen to leverage exclusively the computational chemistry approach described above. While experimental validation of the results herein is of course desirable, they are outside the stated scope of our work, which is strictly computational. By applying molecular dynamics and docking to help elucidate interactions between native and mirror proteins with antibiotic ligands, we have built on the results of previous computational studies. Other groups have studied the interaction of penicillin binding protein 3 (PBP3) from *Chlamydia trachomatis* with amoxicillin.^12^ Our first aim, therefore, was to reproduce the results of these experiments with the exact same simulation setup: ligand, protein model, force field, etc. This provided a baseline of confidence in our study methodology and in the findings of previous work.

Doing so allowed us to meet our second aim: to determine whether the conclusions of previously published academic studies were generalisable across antibiotics with differing physicochemical characteristics. We examined this question by extending a well-established computational methodology to a novel range of antibiotics and bacterial protein targets (Fig. 1). This provides novel, robust *in silico* data that motivates further research into the efficacy of existing countermeasures against hypothetical mirror organisms by the field at large – including through future experimental work that may mirror existing laboratory-based approaches.^10^

**Figure 1.**
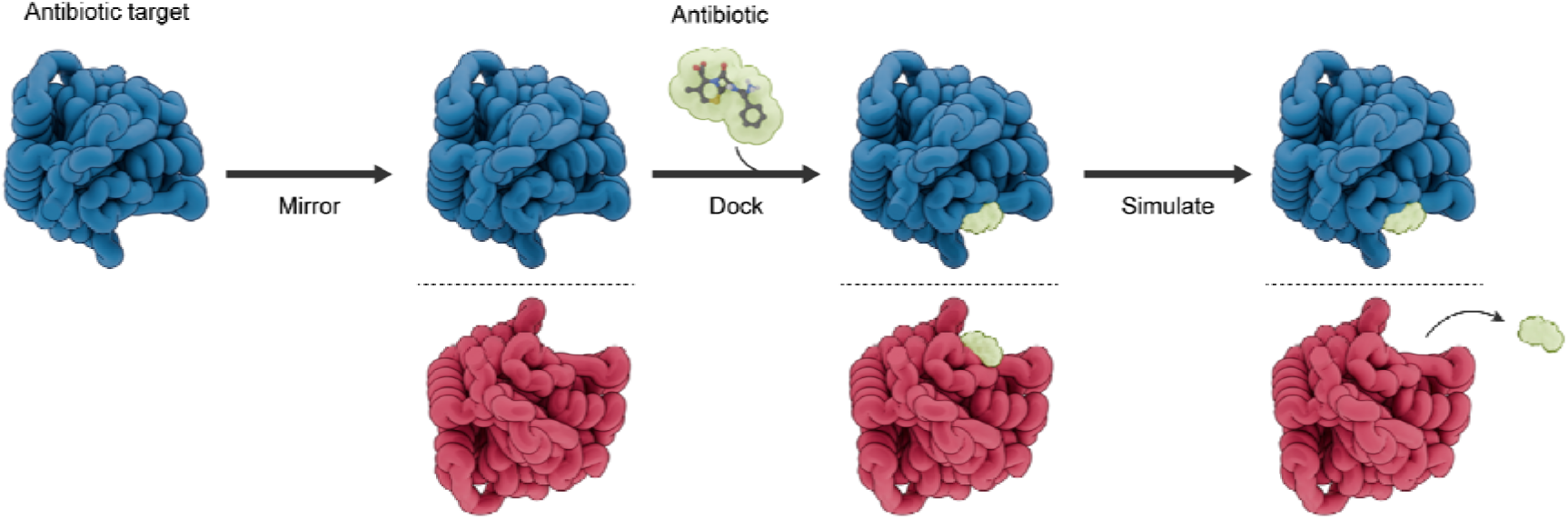
Computational chemistry enables detailed investigations of interactions of antibiotics and their chiral targets, native and mirrored. Rendered schematic depicting simulation workflow, a selection of antibiotic target proteins from pathogenic species were computationally mirrored, to emulate proteins assembled from D-amino acids in mirror life forms. The native, *i.e.* L-amino acids, (navy) and mirror, *i.e.* D-amino acids (scarlet) proteins were simulated at equilibrium, before performing molecular docking with their constituent antibiotics. Further unrestrained molecular dynamics were performed to determine if the antibiotic would remain in its binding site, suggesting it could inactivate the target protein, and function as an antimicrobial.

## 2. Materials and Methods

### Experimental Procedure

Pre-equilibrated antibiotic protein targets and their respective mirror images were simulated in the presence of their antibiotic ligand, using continued ligand proximity as a proxy for target inhibition. High resolution structural data for antibiotic targets were gathered from the Protein Data Bank (PDB) and computationally mirrored. Both mirror and native proteins were then computationally equilibrated, after which their constituent antibiotic was inserted into the binding site via molecular docking. Subsequently, proteins were further simulated in the presence of the antibiotic ligand to compare the potential relative affinity of the antibiotics for the native and mirror form targets.

### Simulation Preparation

Protein targets of antibiotics and their mirror images were first simulated at equilibrium. All-atom protein files for a selection of antibiotic targets were downloaded from the PDB.^13^ The proteins investigated were UDP-*N*-Acetylglucosamine 1-Carboxyvinyltransferase from *E. cloacae* (MurA, PDB:3lth)^14^, Dihydropteroate Synthase from *S. aureus* (DHPS, PDB:1ad1)^15^, the transpeptidase domain of Penicillin Binding Protein 3 from *S. aureus* (PBP3, PDB:6i1f)^16^, and Isoleucyl-tRNA Synthetase from *T. thermophilus* (IleRS, PDB:1jzs)^17^. Mirror image protein structures were generated using a Python protocol developed by Pedroni *et al*.^11,12^ The script applies a full spatial inversion through the origin for every ATOM and HETATM coordinate in a given PDB file, e.g. (x,y,z) becomes (-x,-y,-z), producing a mirror image of the structure. As a result, all stereocenters are inverted and the “handedness” of the molecule is reversed. This converts the naturally occurring L-amino acid configurations (typically S for residues except cysteine) to the corresponding D-amino acid configuration.^18^ This transformation is applied to only the molecular coordinates and not the topology, so atomic connectivity remains unchanged. Inversion of the desired coordinates was verified via inspection of the outputted .pdb files, as well as subsequent visualisation, see SI Fig. 6 for a demonstration of the stereochemical inversion. Structures were then solvated in SPC/E water and neutralised with Na^+^ and Cl^−^ions using *GROMACS* tools (version 2024.3).^19^ Structures were minimised to avoid steric clashes using a steepest descent algorithm for a maximum of 10,000 steps. Systems were then equilibrated to 300 K for 100 ps, using a velocity rescaling thermostat with a 2 ps coupling time. This was followed by isobaric equilibration to 1 bar using isotropic Parrinello-Rahman pressure coupling for 100 ps, also with a 2 ps coupling time (SI Fig. 4). Production simulations were performed for at least 100 ns with at; east two repeats. The CHARMM36 force field^20^ was used and electrostatic forces were calculated using the particle-mesh Ewald (PME) algorithm, and a smooth switching algorithm with a 1.2 nm cutoff was used for van der Waals and electrostatic interactions.

### Molecular Docking

Equilibrated proteins were then docked with their respective antibiotics. Equilibration of native and mirror proteins was assessed by stabilisation of the c-alpha root mean squared deviation (RMSD) and radius of gyration (RoG). Once equilibrated, representative protein structures were prepared for docking, then docked with antibiotic ligands. Protein structures had been relieved of any crystallographic components, e.g. waters or co-crystallised ligands prior to equilibration. To prepare for docking, any non-protein components of the equilibration simulations, such as water or ion atoms were removed. Ionisable amino-acid side chains were maintained from the equilibration, matching physiological conditions at pH 7.5. Protein receptors were then converted to PDBQT format using Meeko^22^, with polar hydrogens and Gasteiger partial charges added. Amino acid side chains were treated as rigid during docking, as the proteins were already equilibrated, and the ligand binding pockets were previously identified. Three-dimensional molecular conformations for the ligands were downloaded from PubChem.^21^ Ligands used were sulfamethoxazole (SMX, CID:5329), fosfomycin (FFQ, CID:446987), amoxicillin (AMX, CID:33613), cephalexin (CPX, CID:27447), and mupirocin (MRC, CID:446596). Ligand tautomers were also protonated at pH 7.5 and converted to PDBQT format with Gasteiger charges ready for docking using Meeko.^22^ Rotatable bonds in the ligands were retained as flexible during the docking. Mirror ligands and mirror proteins were treated identically to their native counterparts to ensure equivalent treatment of stereochemistry, protonation and charge assessment during docking calculations. Antibiotic binding sites were identified using the bound antibiotics from the crystallographic data. In the case of DHPS, where there was no bound ligand, a homologue with the correctly bound ligand was used (PDB:3tzf)^23^ and structures aligned using TM-align.^24^ The docking grid-box was centred on the crystallographic ligand centroid. Grid dimensions were generated automatically using the Meeko receptor preparation utility with a 1.5 Å padding in each dimension around the reference ligand coordinates. Docking was then performed using *AutoDock Vina,*^25^ using a random seed, exhaustiveness was set to 8, the energy range at 3 kcal/mol and the number of modes inspected was 10.^26^ Prior to docking experiments, re-docking of the crystallographic ligands against the antibiotic target proteins was performed to validate the docking procedure, revealing an average heavy atom RMSD of 0.22 ± 0.018 nm.

### Ligand Simulations

Docking poses with the lowest energy minima were taken forward for ligand-bound simulations. Ligand parameters were generated using *CGenFF*^27^ implemented in CHARMM-GUI. All parameters had penalty scores <10, suggesting high analogy to existing parameters, and further optimisation was not required. System topologies were generated using *Gromologist,*^28^ before being prepared, equilibrated, and simulated as described above. Ligand bound trajectories were again simulated for at least 100 ns with a minimum of 2 repeats.

### Analysis

Analysis was performed using *GROMACS* tools. To reduce the impact of equilibration effects, the initial 70% of the trajectories were discarded when performing time-averaged analysis such as RMSD or RoG analysis. *PLIP* was used to identify key binding residues;^29^ *LigPlot+* was used to generate two-dimensional residue interaction maps;^30^ and renders were produced using *VMD*^31^ and Blender. Data analysis and plotting was performed using *R* and *ggplot2*,^32,33^ including the packages *peptides* and *ggradar, bio3d* and *fmsb.*^34–37^

**Figure 2.**
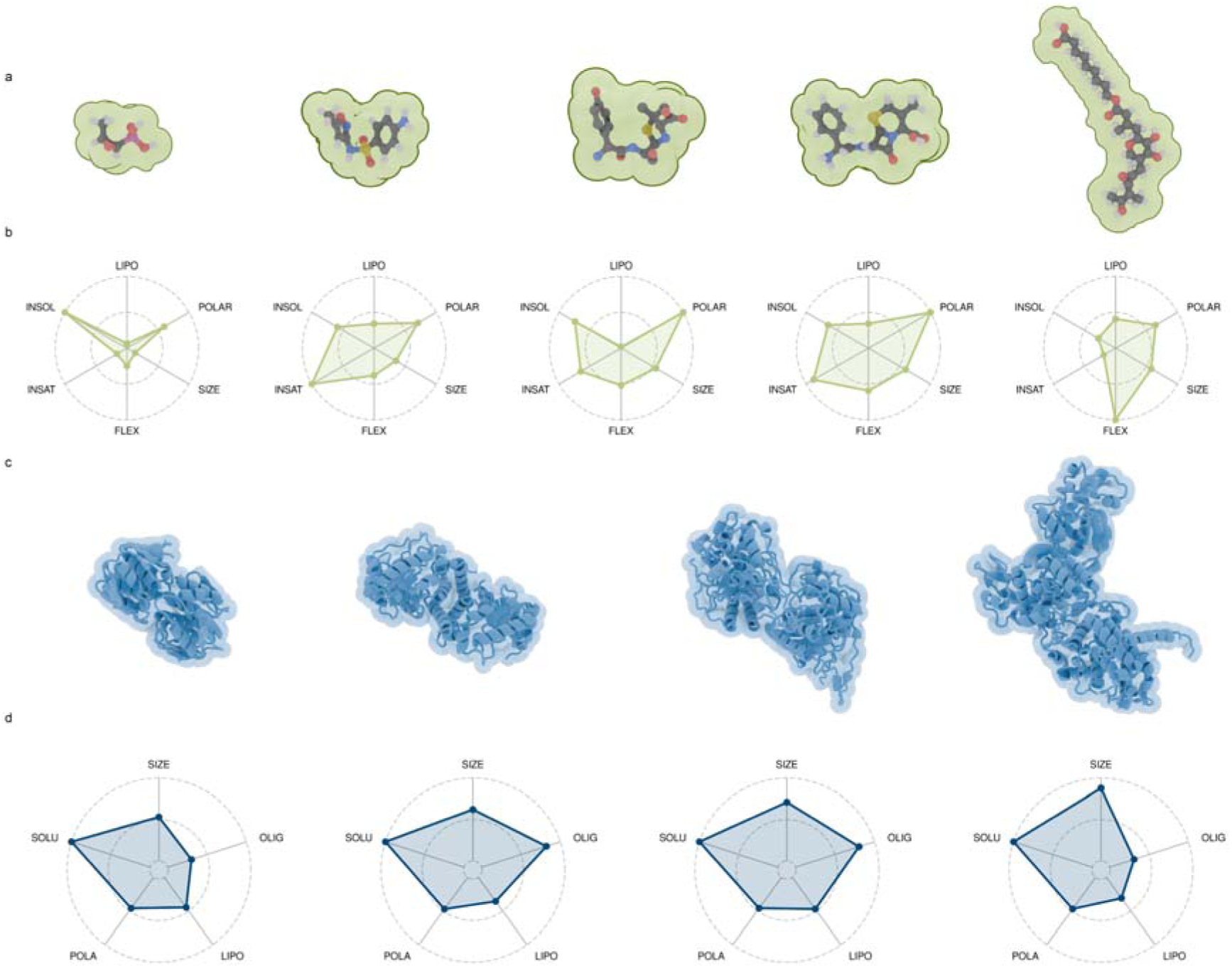
Antibiotics and targets of differing physicochemical and structural properties that were investigated in this work. **(a)** 3D molecular conformers (carbon atoms are depicted grey, hydrogen white, oxygen red, sulphur yellow and nitrogen blue, electrostatic shells are indicated in green) and **(b)** physicochemical properties of antibiotics used in this study, as assessed by SwissADME.^38^ Parameters are lipophilicity (XLogP),^39^ polarity (TPSA),^40^ size (molecular weight), insolubility (ESOL),^41^ insaturation (fraction Csp3) and flexibility (number of rotatable bonds). Left to right shows fosfomycin (FFQ, CID:446987), sulfamethoxazole (SMX, CID:5329), amoxicillin (AMX, CID:33613), cephalexin (CPX, CID:27447), and mupirocin (MRC, CID:446596). **(c)** Space-filling and cartoon structures for the 4 proteins investigated in this study. Left to right is UDP-N-Acetylglucosamine 1-Carboxyvinyltransferase from *E. cloacae* (MurA, PDB:3lth)^14^, Dihydropteroate Synthase from *S. aureus* (DHPS, PDB:1ad1)^15^, the transpeptidase domain of Penicillin Binding Protein 3 from *S. aureus* (PBP3, PDB:6i1f)^16^, and Isoleucyl-tRNA Synthetase from *T. thermophilus* (IleRS, PDB:1jzs)^17^. **(d)** Structural properties of the antibiotic protein targets studied. The molecular weight (SIZE) in kiloDaltons (kDa), normalised between 10 and 240 kDa.^42^ The number of protein chains (OLIG). The exterior lipophilicity (LIPO), normalised between 0 and 1,^43^ calculated using the Wimley-White method.^44^ The average solvent accessible residue polarity (POLA), scaled to between 0 and 80.^45^ The protein solubility (SOLU) calculated as the inverse Boman index,^46^ normalised between 0 and 4.

## 3. Results

### Enolpyruvyl Transferase (MurA)

Enolpyruvyl transferase (MurA) is the smallest enzyme tested; it acts to catalyse the first step in bacterial cell wall biosynthesis.^47^ MurA is a UDP-*N*-acetylglucosamine enolpyruvyl transferase, which generates UDP-GlcNAc-enolpyruvate and inorganic phosphate from phosphoenolpyruvate (PEP) and UDP-N-acetylglucosamine.^48^ The MurA investigated in this work is a 45.58 kDa monomeric protein of 2 globular domains from the Gram-negative bacteria *Enterobacter cloacae*. This species is one of the most common *Enterobacteriaceae,* and acts as an opportunistic pathogen.^49^ *Enterobacteriaceae* infections can result in bacteraemia, sepsis, meningitis, endocarditis and osteomyelitis as well as infections in the urinary and respiratory tracts.^50–52^

**Figure 3.**
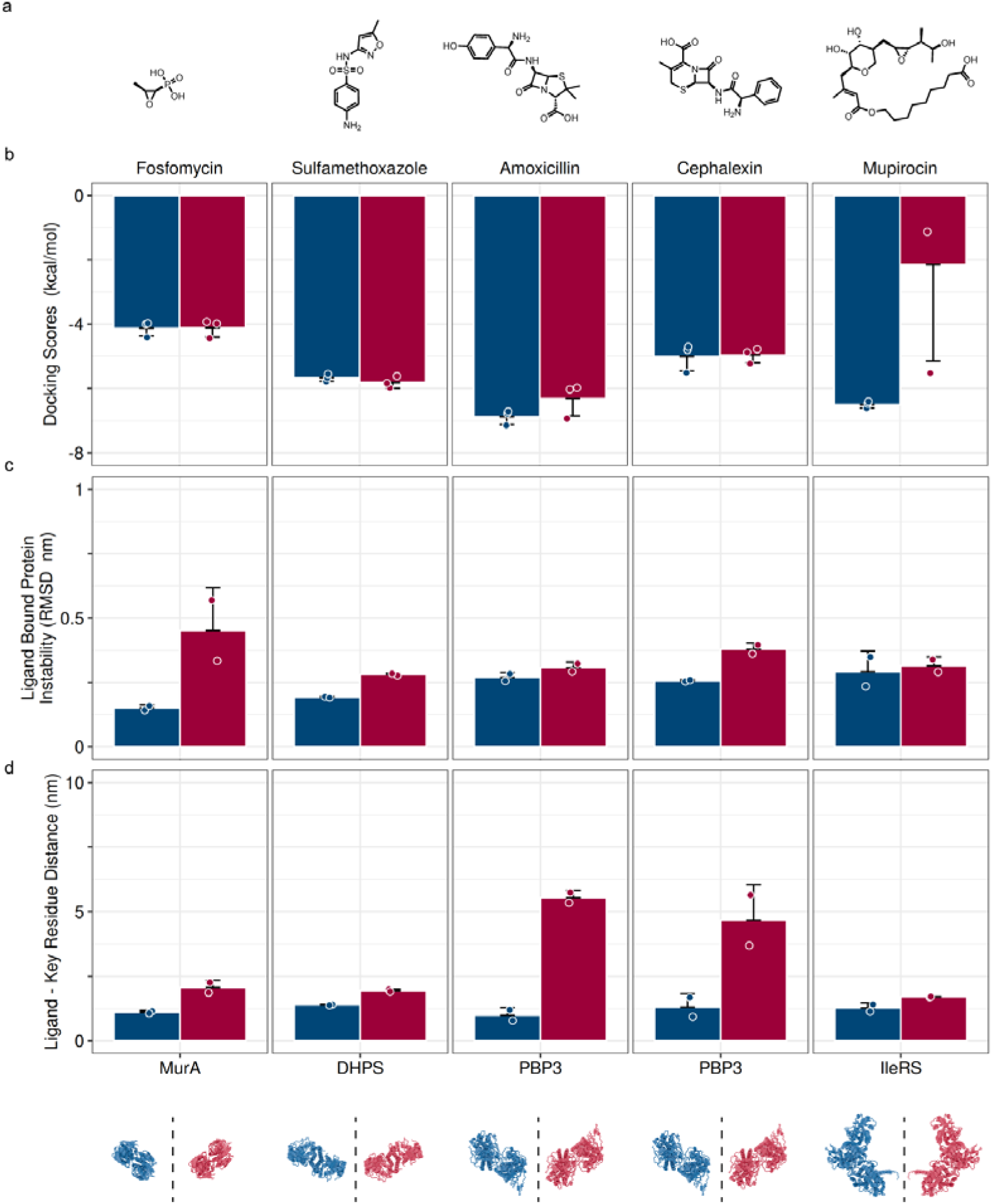
Antibiotics are not effective against mirror targets when compared to their native states, even when well fitted inside the target binding pocket. **(a)** Chemical structures for the antibiotics investigated. **(b)** Docking scores derived when docking antibiotics into the equilibrated native (navy) and mirror (scarlet) targets. **(c)** Protein stability, as measured by alpha carbon RMSD, when simulated after docking with the bound relevant antibiotic. **(d)** Distance between the antibiotic ligand and the key-residue involved in antibiotic inactivation. Each graph shows the average of at least two repeats, the error bars show the standard deviation and the points show the individual data. The data shown here was time-averaged over the terminal 30% of each trajectory. Key residues, determined using PLIP and LigPlot+, as well as literature data are; MurA-fosfomycin, Cys115; DHPS-sulfamethoxazole, Thr51; PBP3-amoxicillin, Ser320; PBP3-cephalexin, Ser320; IleRS-mupirocin, Leu583.

When simulated in solution, both the native and mirror forms of MurA rapidly equilibrated. Equilibration status was determined by assessing the RMSD and RoG values of alpha carbons in the protein. Both the native and mirror forms appeared geometrically stable after the 100 ns equilibration, consistent with previous reports of simulating proteins this size from crystallography data.^53,54^ After equilibration, native MurA reported an RMSD value of 0.16 ± 0.002, while the mirror form reported an increased 0.35 ± 0.1 nm. Equilibration of the structure was further confirmed by the RoG analysis, which rapidly stabilised to 2.23 ± 0.004 and 2.31 ± 0.02 nm (SI Fig. 1).

MurA is inhibited by the action of the antibiotic fosfomycin. Fosfomycin is a 1.38 kDa broad spectrum antibiotic. The antibiotic acts as a competitive inhibitor, entering the PEP binding site and alkylating Cys115.^55^ Cys115 is located within a highly mobile loop within the enzyme’s active site,^56^ and is an essential component in the catalytic action of MurA^57^ and conserved in all bacterial homologues^58^, rendering it an excellent antibiotic target. Alkylation of Cys115 also forces the enzyme into an inactive, closed conformation, irreversibly blocking peptidoglycan synthesis, which is essential for bacterial survival.^59^

Docking of fosfomycin into the active site of MurA was remarkably similar in the native and mirror forms. In both cases the antibiotic fit into a similar groove in the active site, likely due to the flexibility of the Cys115-containing loops, demonstrated by the comparable docking scores of −4.13 ± 0.24 and −4.12 ± 0.28 kcal/mol (Fig. 3b) for the native and mirror form respectively, which are in-line with previous QM/MM calculations of fosfomycin modifying enzymes.^60^

**Figure 4.**
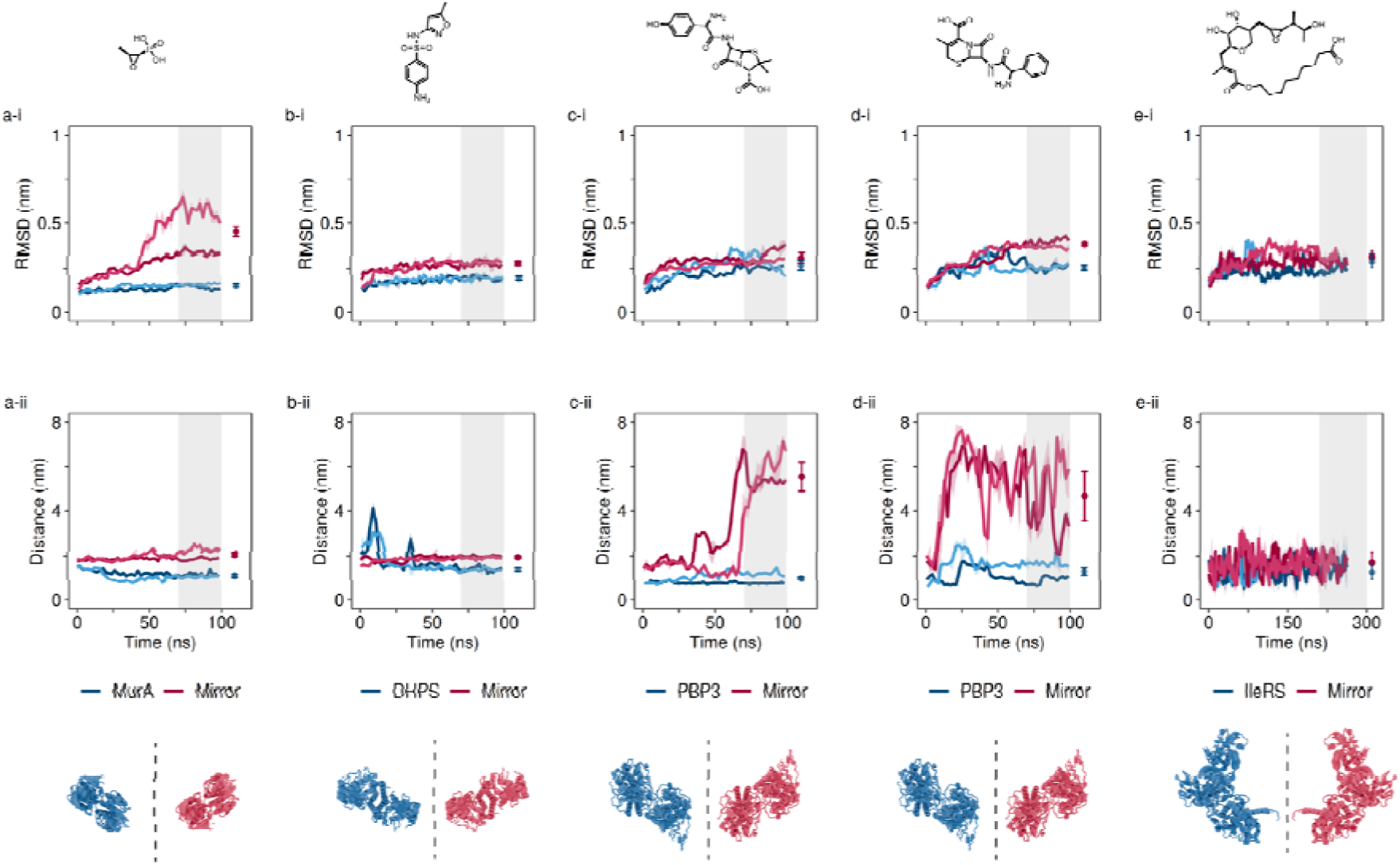
Stability and key residue interactions of antibiotics with their targets and mirrors are antibiotic and target dependent. **(a)** Chemical structure of fosfomycin, **(a-i)** time-averaged alpha carbon RMSD for MurA (navy) and the mirror form (scarlet) once docked with fosfomycin. **(a-ii)** distance between fosfomycin and its key target residue. **(b, b-i, b-ii)** As above, for DHPS and sulfamethoxazole. **(c, c-i, c-ii)** PBP3 and amoxicillin. **(d, d-i, d-ii)** PBP3 and cephalexin. **(e, e-i, e-ii)** IleRS and mupirocin. Each trace is the average of at least two repeats, with the transparent ribbon showing the standard deviation. The grey highlighted region demonstrates the terminal 30% time-period that was used for averaging the final results, and the average and standard deviation for each repeat are shown as points with error on the right-hand side of each plot.

Once the antibiotic-bound forms of MurA were freely simulated, both native and mirror forms were able to retain the antibiotic. While fosfomycin was retained close to the binding site of the mirror form MurA (SI Vid. 2), in both simulations the antibiotic rapidly moved away from the target Cys115 residue (Fig. 4a-ii), with an average distance of 2.06 ± 0.27, compared to 1.11 ± 0.06 nm in the native form (Fig. 3d). In nature, fosfomycin binding is dependent on the initial binding of UDP-GlcNAc to MurA, causing a conformational shift to enable PEP (or its antibiotic analogue fosfomycin) to bind.^61^ In this investigation, this did not appear to be necessary given the tight binding of the native protein to the antibiotic that we observed. Unlike the tight binding of the native form, the distance between the antibiotic fosfomycin and its target Cys115 was large enough that the covalent-bond-mediated inhibition by fosfomycin should not be able to occur, suggesting that the antibiotic may not be effective against mirror form MurA. However, in some simulations the action of fosfomycin caused a significant destabilisation of MurA (Fig. 5, SI Vid. 2). This destabilisation is demonstrated by the average ligand-bound RMSD; 0.15 ± 0.01 for the native vs 0.45 ± 0.17 nm for the mirror. This ligand induced destabilisation may also inhibit the enzyme’s function – but stands in stark contrast to the native action of the antibiotic which induces a closed and non-functional state of the enzyme.

**Figure 5.**
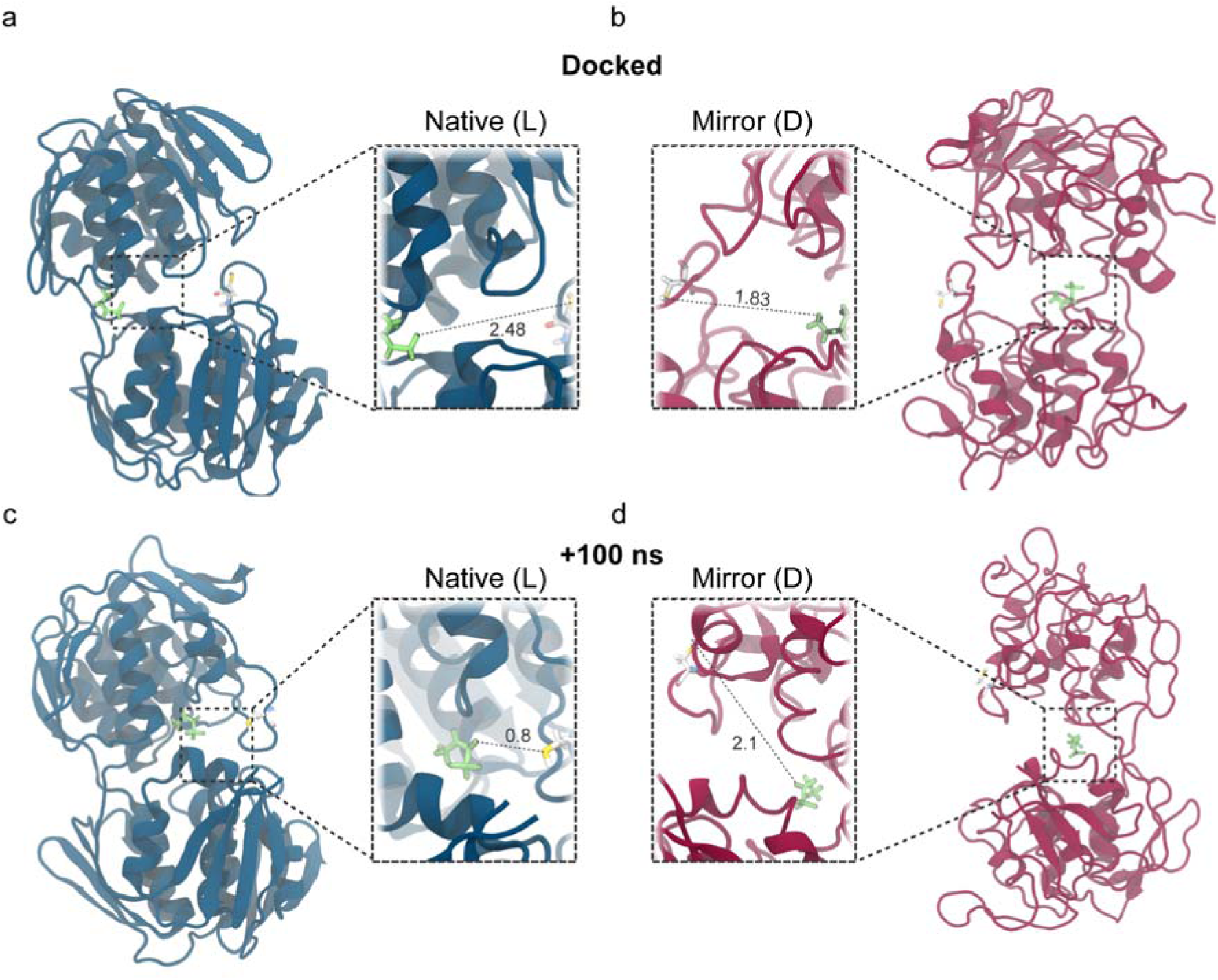
Mirror-MurA retains fosfomycin, but cannot maintain a stable structure. **(a)** Equilibrated native MurA and **(b)** mirror MurA (e.g. D-form) docked with fosfomycin, with Cys 115 highlighted. **(c)** Native and **(d)** mirror MurA after 100 ns of unrestrained dynamics with fosfomycin. Proteins are shown in the cartoon representation, native proteins are navy and mirror forms are scarlet. Both antibiotics and key binding residues are depicted as sticks, all of the antibiotic molecules are shown in lime green, while the key binding residue is shown with carbon atoms in white, oxygen in red, nitrogen in blue, sulphur in yellow and phosphorus in brown. Each zoomed inset shows the current distance between the closest part of the antibiotic and the side-chain of the key binding residue, in nm.

### Dihydropteroate Synthase (DHPS)

Dihydropteroate synthase (DHPS) is an enzyme involved in folate biosynthesis, essential in DNA synthesis and repair.^62^ Folate synthesis is a pathway absent in humans, making DHPS a tractable antibacterial target exploited by a range of existing antibiotics. Here we investigated DHPS from *Staphylococcus aureus,*^63^ a human pathogen involved in a range of diseases including pneumonia, sepsis and osteomyelitis, with a well-characterised and extensive antibiotic resistance profile. DHPS is a dimeric protein with two active sites, with flexible loop regions that contain antibiotic target residues Thr51, Phe17 and Ser18, which act to catalyse the condensation of 6-hydroxymethyl-7,8-dihydropterin-pyrophosphate and *p*-aminobenzoic acid to produce folate.^64^

Equilibration simulations demonstrated rapid plateauing of both native and mirror form DHPS, suggesting geometric stability. Average RMSD values for DHPS in the absence of antibiotics were 0.21 ± 0.01 nm for the native form and 0.29 ± 0.001 nm for the mirror form (SI Fig. 1c). RoG was also used to confirm equilibration, with both forms demonstrating stability with average values of 2.68 ± 0.01 and 2.65 ± 0.008 nm (SI Fig. 1d). The RMSD values we obtained are similar, if slightly higher than previously reported,^65^ likely due to the use of different force fields and omission of stabilising magnesium ions.

DHPS is inhibited by the antibiotic sulfamethoxazole. Sulfamethoxazole is a competitive inhibitor which prevents *p*-aminobenzoic acid from accessing the binding site.^66^ This inhibition is reversible, acting to sterically inhibit the natural substrate as opposed to inactivating key residues. However mutation of key residues such as T51M, which is present in all pathogenic *S. aureus* DHPS variants,^67^ can inhibit sulfamethoxazole binding, while still retaining enzyme function, resulting in clinical resistance.^66^

Docking of sulfamethoxazole into the native and mirror forms of DHPS showed very consistent results. Average docking scores were −5.67 ± 0.12 and −5.81 ± 0.18 kcal/mol (Fig. 3b) for the native and mirror form respectively. In both cases the antibiotic was able to easily enter the active site of the protein. These values are in-line with previous computational work studying the binding of DHPS to other inhibitors, or myoglobin to a sulfamethoxazole analogue, which yielded values between −6.04 kcal/mol and −7.42 kcal/mol respectively.^26,68^

Once bound to sulfamethoxazole, both native and mirror forms showed increased stability over the unbound forms (Fig. 3c). Average RMSD values for the native and mirror forms were 0.19 ± 0.003 and 0.28 ± 0.005 nm respectively, suggesting stabilisation of both forms, although the native state was marginally more stable. In both cases the antibiotic is well retained in the active site, while it is not necessarily always close to the key residue selected (Thr51), this is not necessary for effective inhibition. Average distances between Thr51 and sulfamethoxazole were 1.4 ± 0.02 nm and 1.95 ± 0.05 nm for the native and mirror form respectively. In the native state, some re-organisation of the antibiotic is visible (SI Vid. 3) which is not visible in the mirror form, allowing the antibiotic closer to T51. However it is likely that in both cases the activity of the enzyme would be sterically impeded.

### Penicillin-Binding Protein 3 (PBP3)

Penicillin-Binding Protein 3 (PBP3) is an essential enzyme involved in peptidoglycan crosslinking during cell division.^69^ The PBP3 form tested was from *Chlamydia trachomatis*, an obligate intracellular pathogen commonly involved in urogenital infections, which can result in chronic inflammation and scarring. PBP3 is a 73.67 kDa enzyme, however only the 36.5 kDa TP domain was simulated.^16^ The TP domain was selected both to reduce compute requirements, and to confirm the simulation methodology used here gave results consistent with that of previous work from Pedroni *et al.*^11^

As with the previous proteins, the small TP domain rapidly reached geometric stability. Average RMSD values for PBP3 in the absence of antibiotics were 0.27 ± 0.02 for the native form and 0.31 ± 0.02 for the mirror form (SI Fig. 1e), consistent with the work from Pedroni *et al*. Further demonstration of equilibration came from the stable RoG of the native form (1.98 ± 0.01), which was consistent with that of the mirror form (1.97 ± 0.02), which had fully stabilised after the 100 ns equilibration, despite briefly spiking at 50 ns (SI Fig. 1f).

PBP3 was assessed with 2 different antibiotics, the widely used β lactams amoxicillin and cephalexin. Both antibiotics inhibit cell maturation and differentiation.^70^ The antibiotics form a covalent adduct with Ser320 in the PBP3 active site, inhibiting peptidoglycan cross-linking.^71^ The active site architecture, including the catalytic serine, of PBP is conserved in all PBP3 family proteins^72^ although the residue numbering can differ.^73^ This high degree of conservation in PBP3 proteins, which are essential in Gram-negative bacterial cell division, makes it an attractive target for broad-spectrum β-lactam antibiotics.

Docking PBP3 in complex with amoxicillin and cephalexin showed comparable binding strengths. In all cases the antibiotic could fit into the active site of both the native and mirror form. The average docking scores of the top 3 docking poses were −6.88 ± 0.23 and −6.31 ± 0.54 kcal/mol for the native and mirror PBP3 binding to amoxicillin, and −5.01 ± 0.44 and −4.96 ± 0.24 kcal/mol for cephalexin (Fig. 3b). Experimental data for amoxicillin and cephalexin binding to PBP3 reveal IC_50_ values of 3 μg/mL and 30 μg/mL respectively, which would translate to binding affinities of −7.3 kcal/mol and −5.8 kcal/mol at 25 °C, in-line with the simulated values.^74^

When simulated with the docked antibiotics, both the native and mirror proteins were stable, but the mirror form was unable to retain the antibiotics in close proximity. Protein stability was consistent across repeat simulations with RMSD values of 0.27 ± 0.02 and 0.31 ± 0.02 nm for the native and mirror forms complexed to amoxicillin (respectively) and 0.26 ± 0.004 and 0.38 ± 0.02 nm for the native and mirror forms complexed to cephalexin respectively (Fig. 3c). Despite this stability, the distance between the antibiotics and their target residue Ser320 demonstrated the ineffectiveness of the antibiotics against the mirror form bacterial target. In the native forms amoxicillin was retained within 0.99 ± 0.3 nm of Ser320, while cephalexin was 1.31 ± 0.52 nm (Fig. 4c-ii, 4d-ii). However, the mirror forms saw the antibiotics exit the binding site within 10 ns for cephalexin and 60 ns for amoxicillin, with final distances being 5.54 ± 0.3 nm for amoxicillin and 4.67 ± 1.4 nm for cephalexin (Fig. 4c-ii, 4d-ii). This strongly suggests that the antibiotics would be ineffective – although it may be feasible that amoxicillin could have some long-lasting inhibitory action in the 60 ns it was retained in the binding site, this is a very short timeframe in which to act and it is evidently not well retained compared to the native form of the protein (Fig. 6).

**Figure 6.**
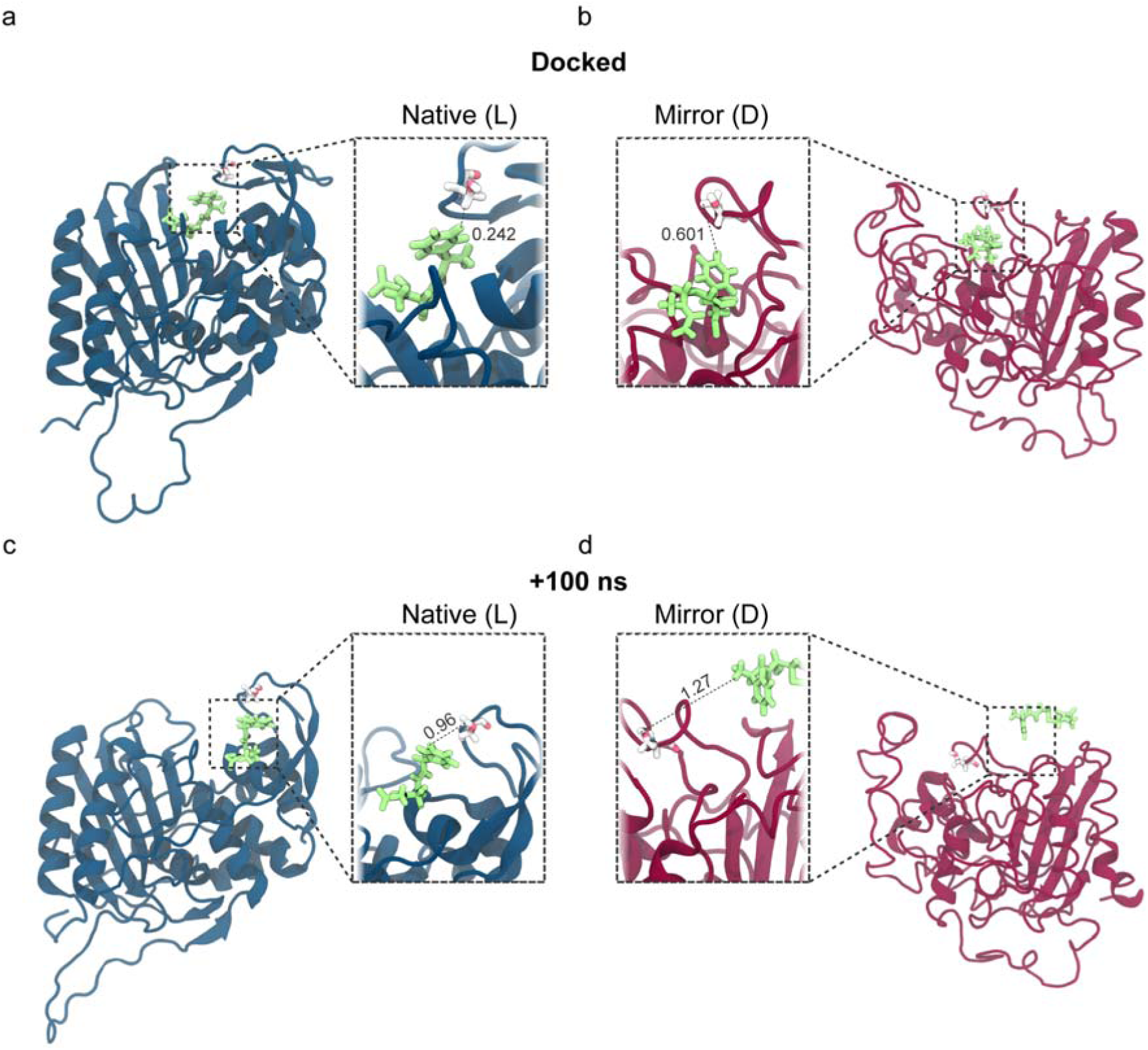
Mirror PBP3 cannot retain cephalexin within the binding pocket. **(a)** Equilibrated native PBP3 and **(b)** mirror PBP3 (e.g. D-form) docked with cephalexin, with Ser320 highlighted. **(c)** Native and **(d)** mirror PBP3 after 100 ns of unrestrained dynamics with cephalexin. Proteins are shown in the cartoon representation, native proteins are navy and mirror forms are scarlet. Both antibiotics and key binding residues are depicted as sticks, all of the antibiotic molecules are shown in lime green, while the key binding residue is shown with carbon atoms in white, oxygen in red, nitrogen in blue, sulphur in yellow and phosphorus in brown. Each zoomed inset shows the current distance between the closest part of the antibiotic and the side-chain of the key binding residue, in nm.

The computational process used to generate the mirror proteins could also be performed on the antibiotics, so a chiral mirrored form of amoxicillin was assessed against native and mirror PBP3. When tested our results were comparable with previous work,^12^ where the mirrored antibiotic could be retained in the mirrored protein. Building on previous work we also investigated the efficacy of the mirror antibiotic on native PBP3, in which predictably the mirror antibiotic was not well retained in the active site of the native protein (SI Fig. 5). The mirrored chirality antibiotic was retained within active site of the mirror PBP3, 1.7 ± 1.9 nm from Ser320, compared to the 2.1 ± 1.3 nm in the native form.

### Isoleucyl-tRNA Synthetase (IleRS)

**I**soleucyl-tRNA synthetase (IleRS) is an essential aminoacyl tRNA synthase enzyme. It acts to assemble isoleucine-tRNA complexes involved in protein synthesis by transferring aminoacyl adenylate to the 3’ terminal adenosine of tRNA.^75^ The IleRS investigated here is from the thermophilic eubacterium, *Thermus thermophilus,* selected for the high quality crystallography data in complex with the antibiotic mupirocin.^17^ While *T. thermophilus* is not a human pathogen, tRNA synthesis is highly conserved and there is no significant difference in the amino acid sequence between catalytic domains of IleRS proteins from different kingdoms.^75^

As the largest protein studied, IleRS showed some of the highest RMSD and RoG values, but still appeared to rapidly reach geometric equilibrium. The 95.31 kDa protein appeared to reach stability in both native and mirror forms within the first 10 ns and RMSD and RoG were consistent from that time onwards. Average RMSD for the native IleRS was 0.29 ± 0.03 nm, and the mirror form again showed slightly higher RMSDat 0.35 ± 0.001 nm (SI Fig. 1g). Despite the small change in stability induced by computational mirroring, the RoG values were consistent with both native and mirror forms showing 3.46 ± 0.002 and 3.46 ± 0.002 nm for the native and mirror respectively (SI Fig. 1h).

IleRS is inhibited by the action of the antibiotic mupirocin. Mupirocin is an analogue of the isoleucyl-adenylate substrate, competitively excluding it from the active site and disabling protein synthesis.^75^ Mupirocin is believed to bind to Leu583 in the active site, as its substitution in eukaryotic IleRS confers resistance to mupirocin.^75,76^ This lack of efficacy against eukaryotic IleRS, and the relatively high degree of conservation of Leu583 in bacterial homologues^77^ make mupirocin an effective antibiotic.

The docking of mupirocin into the active site of IleRS showed the largest difference between the native and mirror state of any of the proteins tested. Native IleRS showed highly consistent docking, with a binding energy of −6.5 ± 0.11 kcal/mol. However the mirrored form was highly inconsistent, the top docking pose had a comparable binding energy to the native (Fig. 3b), but the rest were bound significantly less strongly, resulting in an average binding energy of −2.2 ± 3 kcal/mol. Despite the binding energy being significantly reduced compared to the native protein, it was still spontaneous suggesting binding would occur. The pose with the docking score of −6.5 kcal/mol was carried forwards for further simulation as it is comparable with previously published data for the IleRS-mupirocin binding affinity; −7.6 kcal/mol.^78^

IleRS from *T. thermophilus* is the only protein tested to show comparable binding and stability to its cognate antibiotic between the native and mirrored form. The antibiotic bound trajectories had comparable RMSDs (Fig. 3c) of 0.29 ± 0.08 and 0.31 ± 0.03 nm for the native and mirror states respectively, nearly identical to the unbound RMSD. The average distance between the mupirocin and its target residue, Leu583, was 1.28 ± 0.19 and 1.71 ± 0.1 nm in the native and mirror state respectively (Fig. 3d), with the antibiotic staying tightly bound into the active site in both the mirror and native forms (Fig. 4e-ii, Fig. 7). This may be due to the highly flexible nature of both mupirocin and IleRS. IleRS is highly flexible, a known feature of proteins from thermophilic species (Fig. 2b).^79^ However, the variable docking scores suggest that it may be unlikely for the antibiotic to spontaneously enter the binding site, even if it can be retained once there. It is interesting that the only instance of possible antibiotic effectiveness against both native and mirror forms in our work comes from a species that is not pathogenic to humans. It is possible this effect derives from the extremophilic nature of the protein. Extremophilic proteins tend to show increased local flexibility^80^ (Fig. 2d) rendering ligand binding pockets more tolerant to structural change.^81,82^ In conjunction, the long-carbon chain of mupirocin (Fig. 2b) also confers greater flexibility, which may also enhance to binding tolerance.^83^

**Figure 7.**
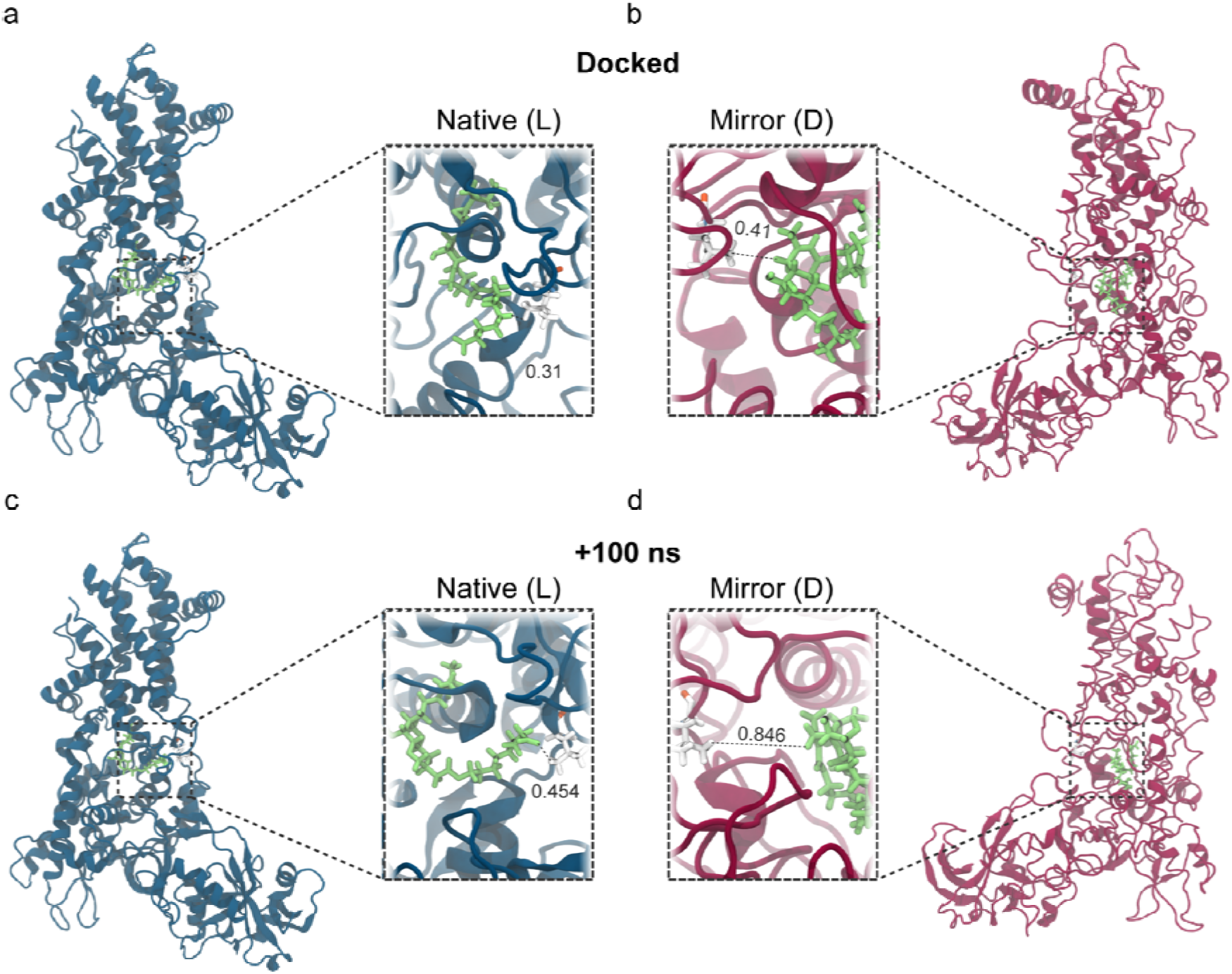
Flexible thermophilic pathogen targets were better able to retain their antibiotic ligand. **(a)** Equilibrated native IleRS and **(b)** mirror IleRS (e.g. D-form) docked with mupirocin, with Leu583 highlighted. **(c)** Native and **(d)** mirror IleRS after 100 ns of unrestrained dynamics with mupirocin. Proteins are shown in the cartoon representation, native proteins are navy and mirror forms are scarlet. Both antibiotics and key binding residues are depicted as sticks, all of the antibiotic molecules are shown in lime green, while the key binding residue is shown with carbon atoms in white, oxygen in red, nitrogen in blue, sulphur in yellow and phosphorus in brown. Each zoomed inset shows the current distance between the closest part of the antibiotic and the side-chain of the key binding residue, in nm.

## 4. Discussion

In most cases, antibiotics were not able to remain in close proximity to binding pockets of mirror forms of bacterial proteins – unlike native forms, despite initially favourable docking. Docking results showed spontaneous interaction for all cases, and in all examples apart from mirrored IleRS, were highly consistent between native and mirror proteins. The docking scores were further supported by MM/PBSA analysis of the ligand-bound trajectories, where the free energy calculations agreed with the docking scores, further suggesting spontaneous binding (SI Fig. 7). It should be noted that the trajectories generated here only sampled the lowest-energy docking poses, and further sampling may reveal more of the energetic landscape. However, the high consistency of docking scores, and the substantial conformational relaxation time for both ligand and receptor reduces the dependence of the conclusions on the initial docking geometry. In many cases, (Fig. 4b-ii), significant spatial re-arrangement occurred until a stable ligand-receptor equilibrium was reached.

From this data, it may be inferred that the antibiotics tested would fail to exert an antimicrobial effect on potential future mirror bacteria. However, it must be noted that these simulations were performed using a non-reactive force field, CHARMM36, which does not explicitly model bond-breaking or bond-formation events. As a result, this data may under-state the effectiveness of antibiotics which rely on covalent modification of their targets, such as amoxicillin. More rigorous approaches using QM/MM simulations of reactive force fields (e.g. ReaxFF) exist, but are associated with increased compute cost and parameterisation challenges, particularly for the large bio-molecular systems studied here. As such, classical force fields are still widely used for studying ligand-protein binding kinetics^84^ and have been well validated experimentally.^85^ Thus, this work provides insight into whether the antibiotic ligands and their target proteins can adopt the stable binding configurations required for subsequent covalent modification steps.

This study has a number of limitations, in particular that it was a small-scale computational study. Longer simulations, of a wider range of antibiotics and protein targets, with more exhaustive conformational sampling or ensemble-based approaches would bolster the findings here. While these were beyond the scope of this exploratory study, we hope that our findings motivate further research by other groups. These simulations evaluate isolated protein–ligand interactions only; whole-cell susceptibility additionally depends on membrane permeability, transporter uptake, efflux, metabolism and cell-envelope architecture, which may differ substantially in mirror bacteria and are not captured here; for instance, sulfamethoxazole efficacy also depends on pathway-level folate metabolism beyond DHPS binding. This limits our ability to infer or draw strong conclusions regarding the ultimate antimicrobial efficacy of existing countermeasures against mirror bacteria. However, our work sits within a small but growing corpus of literature providing preliminary data on interactions between mirror bacterial macromolecules and existing countermeasures,^11,12^ and proxy data on mirror countermeasures and existing bacteria.^10^ While imperfect models, these studies have broadly demonstrated lower binding affinities and efficacies.

Ideally, future work would also include experimental data to validate the relevance of the computational methodology we have adapted and employed. Both the present study and previous work from Pedroni *et al.* form computational mirror systems via coordinate inversion. This forms geometrically valid mirror counterparts, and our data has indicated that it well preserves the structural organisation of the proteins (SI Fig. 2), however as this work is purely computational, it cannot establish that this corresponds to experimentally producible proteins. The CHARMM36 force field has been extensively parameterised and optimised for L-amino acid form proteins, and while bonded interactions are formally invariant, stereochemical inversion here appears to have slightly altered dihedral bond energies (SI Fig. 3) – although within measurement error, likely affecting subsequent side-chain packing geometries, which could have knock-on effects on the hydrogen-bonding networks and solvent organisation of the proteins. This work does not suggest that these changes would arise under *de novo* folding of mirror proteins, and instead focuses on comparative analysis of the mirrored ligand-target interactions within a consistent computational framework. Further work that validates the realism and behaviour of mirror proteins would further establish this precedent, as well as enable optimisation of computational force fields for mirror proteins – but may also act as an enabler in the development of physical mirror life forms and the potential threats that they represent. There are already early attempts to define the efficacy of existing medical countermeasures against mirror bacteria.^10^ Further work of this kind, in addition to *in vitro* binding kinetics experiments between mirror proteins and antibiotics, would help to confirm our findings.

While most antibiotics were shown to be ineffective, antibiotics which act through steric hindrance (as opposed to covalent modification) appear to have greater potential against mirror life, as the conditions for effectiveness are often less spatially stringent. This is likely to depend on the size and flexibility of the binding pocket. This is worrying as antibiotics which work by covalent modification are often more potent and selective antibiotics,^86^ as well as one of the few types of antibiotics in which we continue to see discoveries.^87^ Some of the most historically important and widely used antibiotics, such as the β-lactams, work through covalent modification of target proteins.^88^ Antibiotics which are highly flexible, or target flexible proteins, are likely to be more effective against mirror life. This is a positive effect that warrants further investigation as part of any future anti-mirror drug development efforts. These could draw on existing drug development strategies, as many antibiotics such as linezolid and related oxazolidinones have been optimised by increasing their rotational freedom.^89^ Further, many more modern antibiotics (especially peptide mimics)^90^ are highly flexible and may as a result be more effective against mirror life.

Despite the limitations of exploratory computational data presented here, our findings add to the list of experiments which suggest that existing antibiotics may not be effective against mirror bacteria. The creation of mirror life represents a serious biosecurity threat; the extent to which we would be able to mitigate this threat with existing medical countermeasures warrants significant further investigation.

## Acknowledgments

The authors acknowledge the use of the UCL Kathleen High Performance Computing Facility (Kathleen@UCL) and associated support services in the completion of this work. The authors also acknowledge Ana Leonescu for insightful input regarding the chemistry of small molecule antibiotics.

## Author contributions

Paul-Enguerrand Fady: Conceptualization, Project administration, Writing – original draft, Writing – review & editing, Funding acquisition.

Juliette Ciccone: Investigation, Formal analysis, Data curation, Methodology, Visualization, Writing – original draft, Writing – review & editing.

## Data Availability

All trajectories are available upon request.

## Supplementary Materials

**SI Figure 1.**
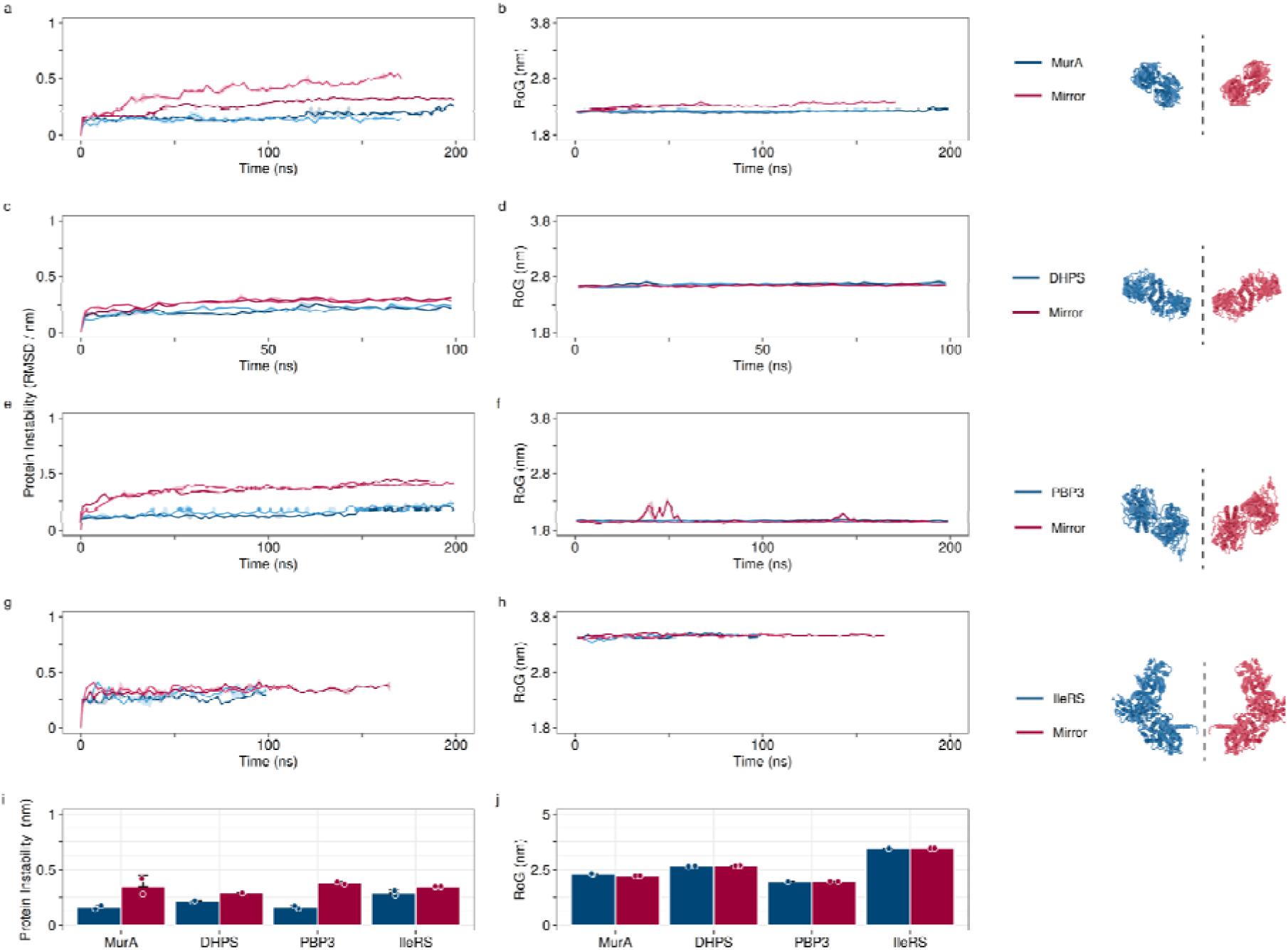
Native and mirror form proteins reached geometric equilibrium after 100 ns. **(a)** Time-averaged alpha carbon RMSD and **(b)** RoG for MurA, **(c-d)** DHPS, **(e-f)** PBP3 and (**g-h)** IleRS. Native proteins are shown in navy and mirror in scarlet. Each trace is an average of 2 repeats and the coloured ribbon shows the standard deviation. **(i)** Average RMSD and **(j)** RoG from the final 30 ns of each trajectory. Each graph shows the average of at least two repeats, the error bars show the standard error of the mean and the points show the raw data.

**SI Figure 2.**
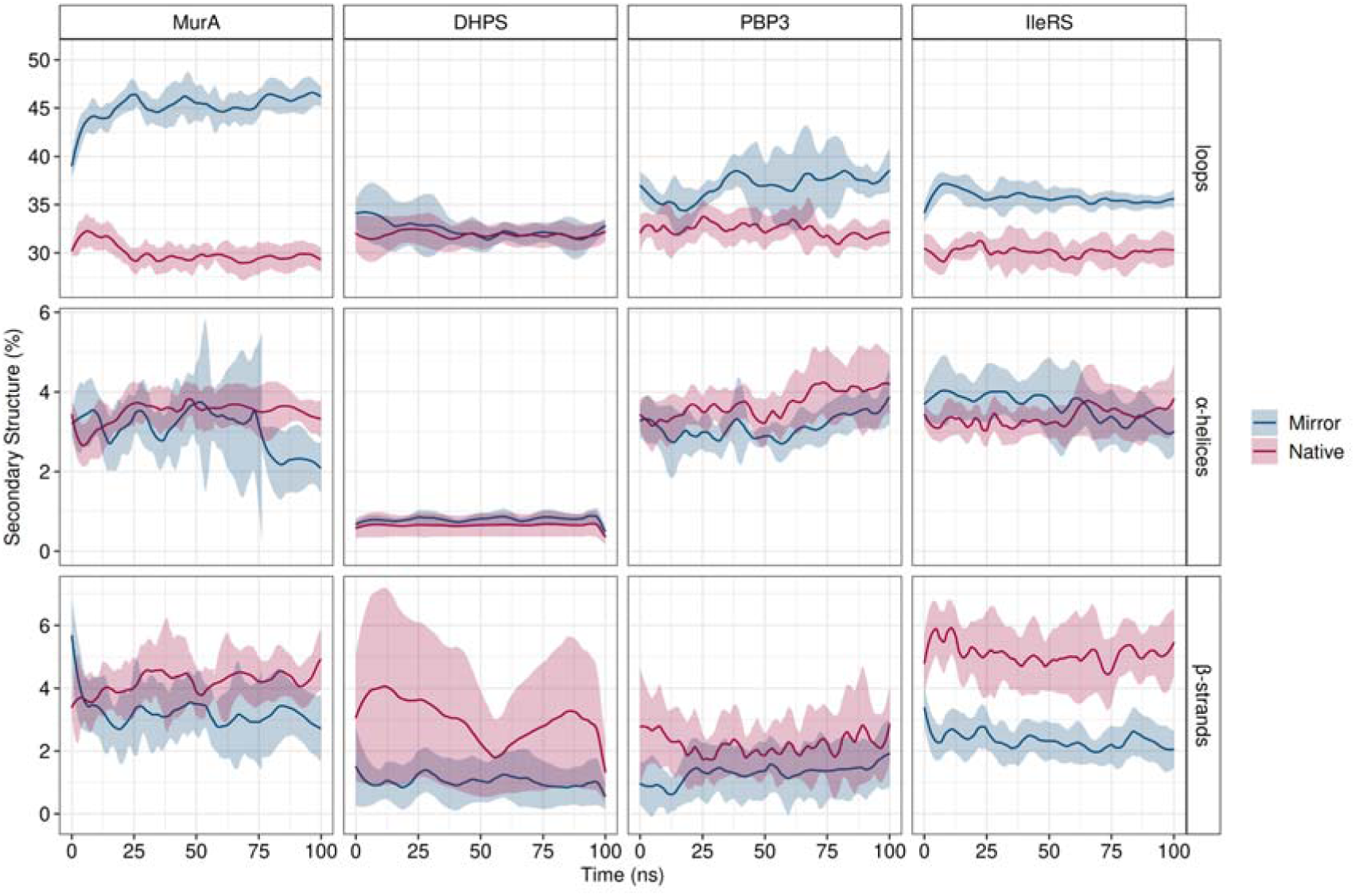
Computational mirroring of proteins retains most secondary structure. Using DSSP (Dictionary of Secondary Structure in Proteins),^91^ the percentage of each protein that exists as a loop, alpha helix or beta strand was calculated for the equilibration simulations. Each trace is an average of two repeats, and the coloured ribbon shows the standard deviation.

**SI Figure 3.**
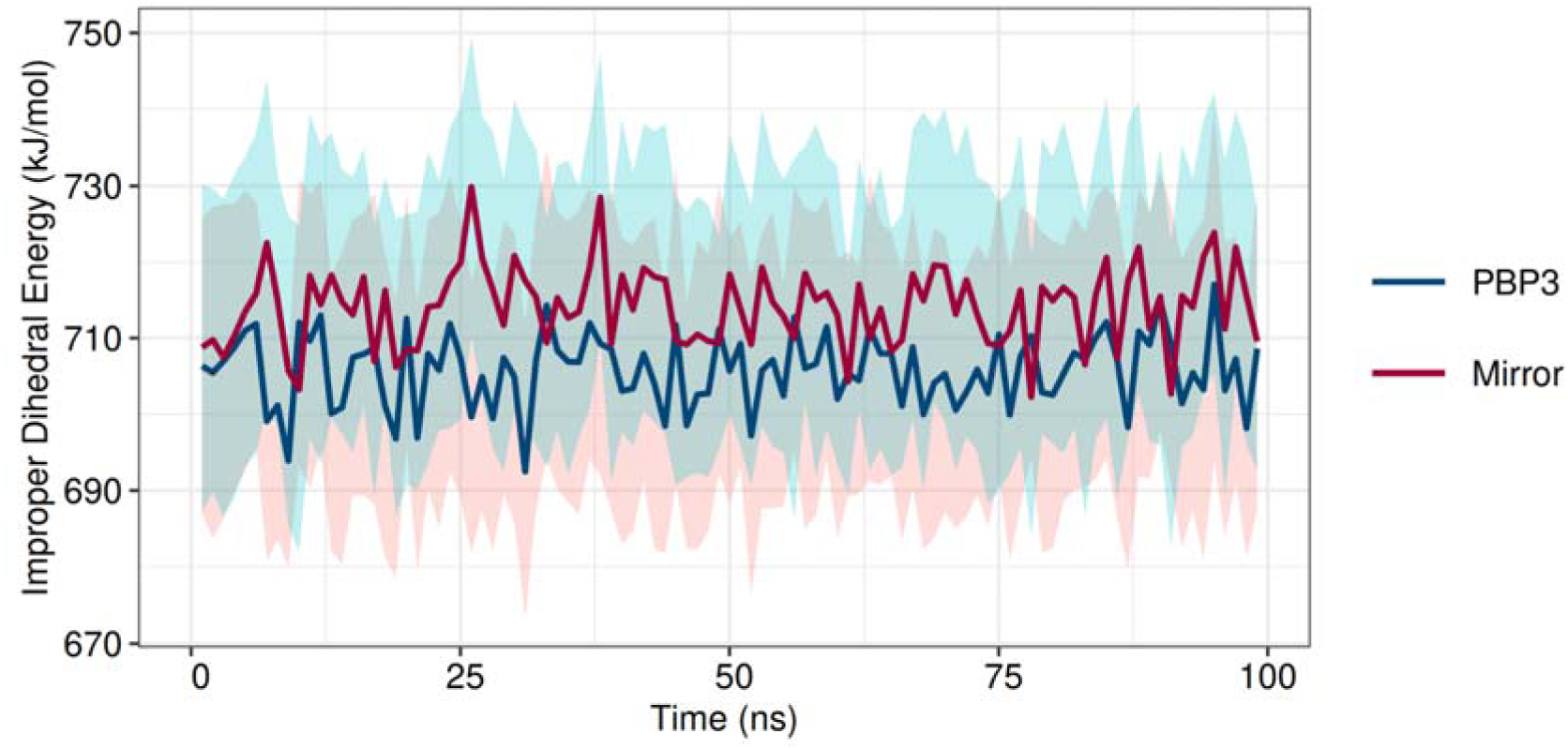
Improper Dihedral Energy for representative protein and mirror counterpart show very little difference. Each line is an average of 2 independent repeats, and the coloured ribbon shows the standard deviation, a 1 ns running average has been applied.

**SI Figure 4.**
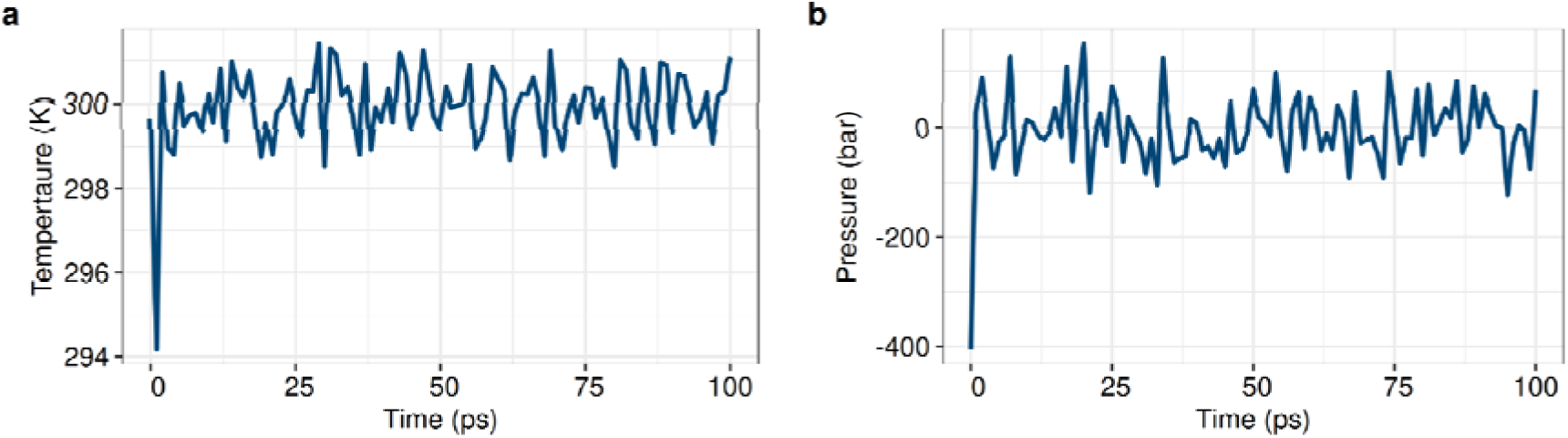
A representative example demonstrating 100 ps is adequate for (a) isothermal (b) 577 isobaric equilibration.

**SI Figure 5.**
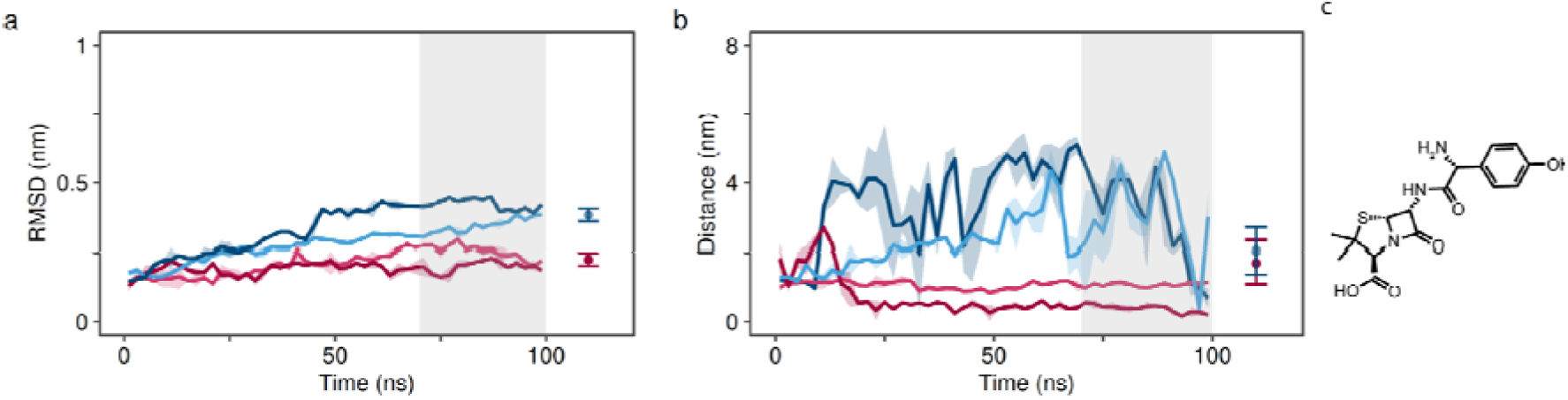
Stability and Ser320 distance for PBP3 when binding to mirrored amoxicillin. **(a)** Alpha carbon RMSD of native PBP3 (blues) and the mirror form (reds) after docking with a mirrored amoxicillin. **(b)** Distance between the centre of mass of amoxicillin and Ser320, a key binding residue in PBP. Each individual trace is shown, with a 2 ns running average, with the coloured ribbon showing the standard deviation. To determine equilibrium values for comparison, the final 30% (shown in grey) of each trajectory was averaged to reveal; RMSD values of 0.34 ± 0.06 and 0.23 ± 0.04 nm, and a centre of mass distance between mirrored amoxicillin and Ser320 of 2.05 ± 1.35 and 1.72 ± 1.9 nm for the native and mirror forms respectively, shown as points to the right of the plots. **(c)** shows the structural formula of the computationally mirrored amoxicillin.

**SI Figure 6.**
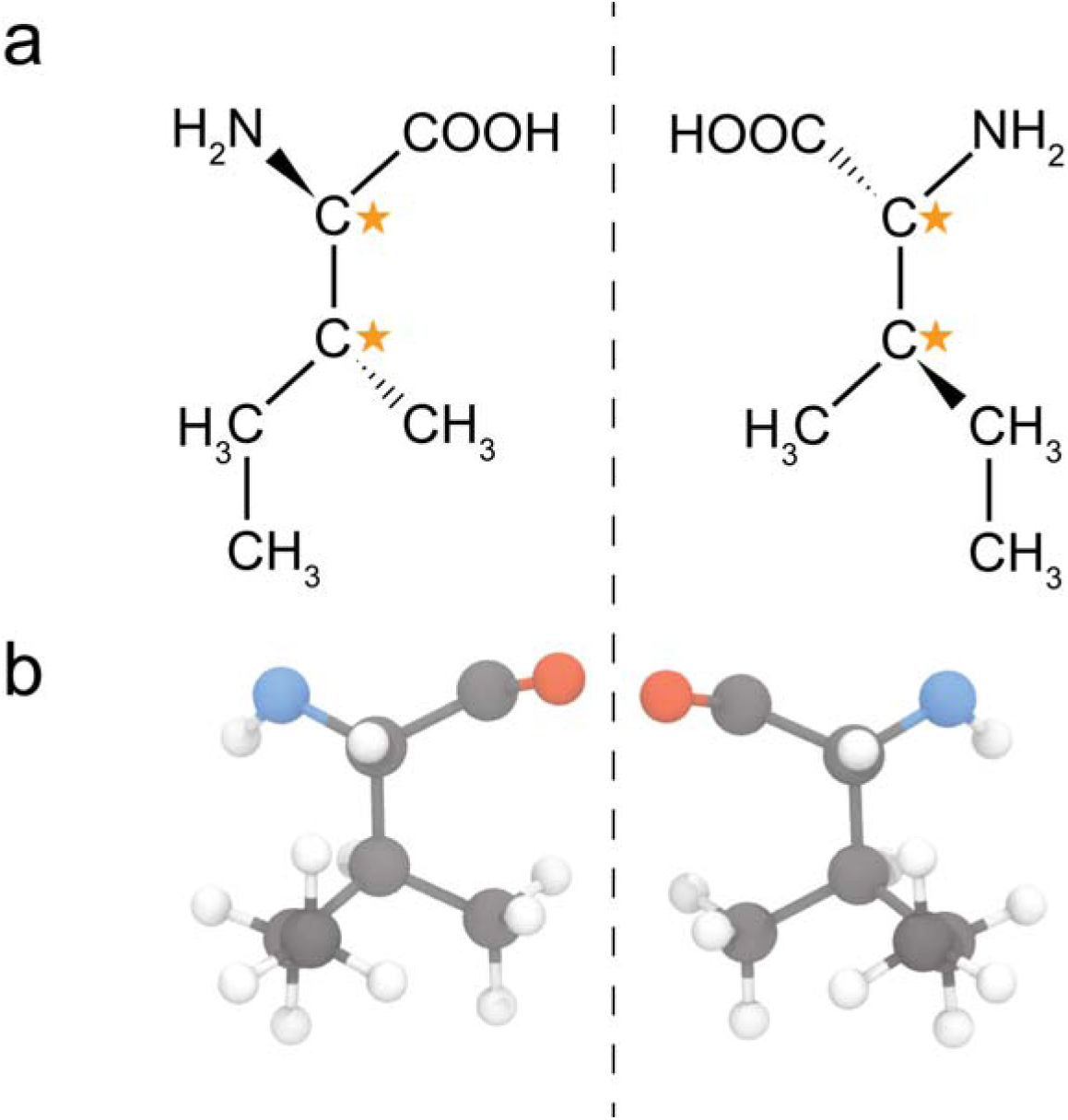
Demonstration of computational chiral inversion. **(a)** Structural formula of an isoleucine amino acid showing the L (native) and D (mirror) forms on the left and right respectively. The chiral centres are indicated with a golden star. **(b)** schematic render of an isoleucine residue that has been through the computational chiral inversion, where the x, y, z coordinates of each atom are converted to −x, −y, −z. Carbon atoms are coloured grey, hydrogen white, nitrogen blue and oxygen red. Note that the structure in b is missing the monomer hydrogens shown in a as it is part of an amino acid polymer.

**SI Figure 7.**
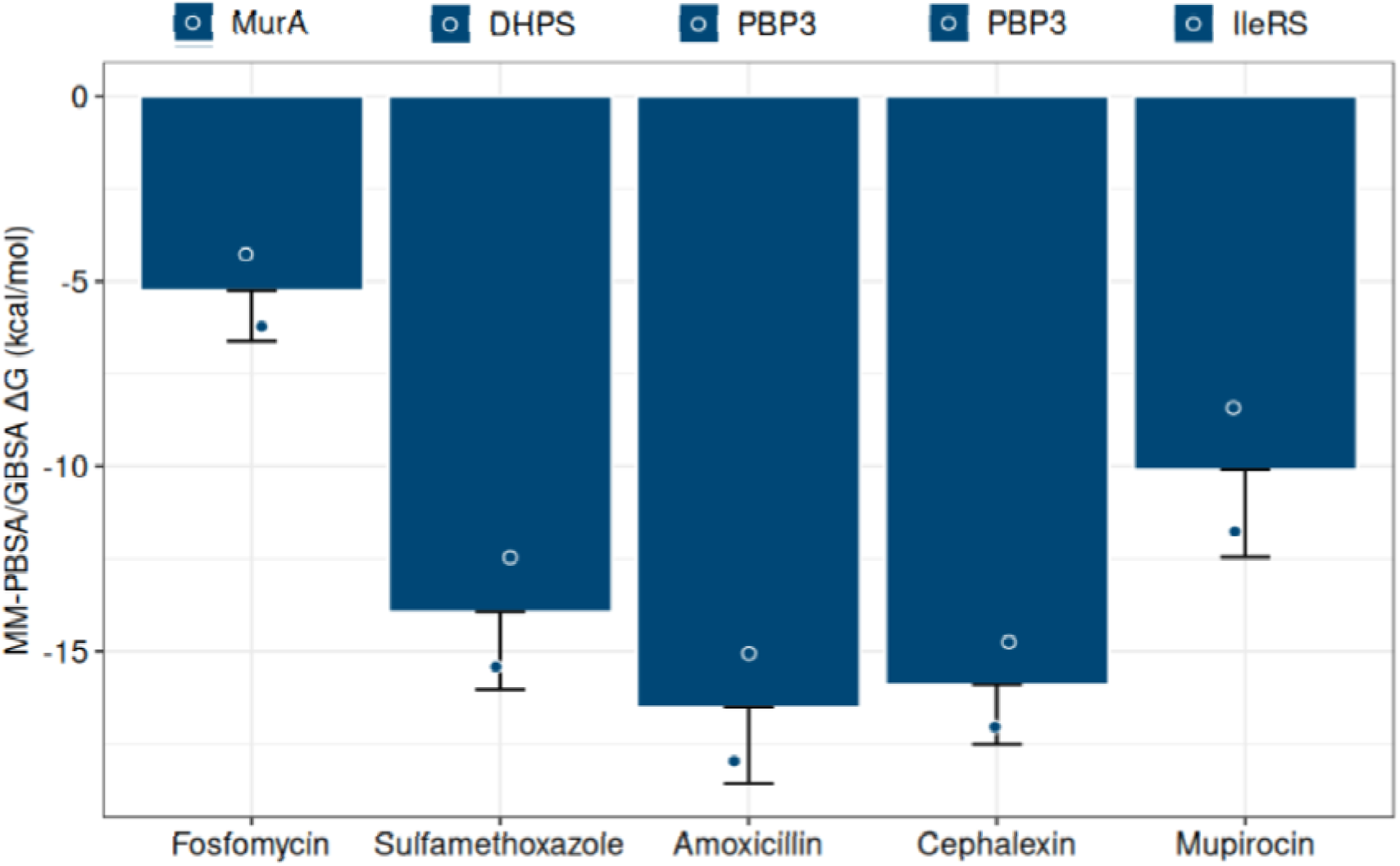
MM/PBSA analysis of docked trajectories agrees with docking scores. Binding free energies were estimated using the MM/PBSA approach implemented in g_mmpbsa and was performed on a 20 ns snapshot at the end of the ligand-bound trajectory of the native form proteins, sampled at 1 ns intervals. A protein dielectric constant of 4 was used throughout, polar solvation energies were obtained by numerical solution of the Poisson–Boltzmann equation, while non-polar contributions were calculated from solvent-accessible surface area terms. Average scores are shown as bars, with individual data as points and the standard deviation shown as error bars.

**SI Table 1-.** Root Mean Squared Deviation.

| Protein | Ligand | Enantiomer | RMSD (nm) | StDev |
| --- | --- | --- | --- | --- |
| MurA | Unbound | Native | 0.16 | 0.02 |
| MurA | Unbound | Mirror | 0.35 | 0.1 |
| DHPS | Unbound | Native | 0.21 | 0.009 |
| DHPS | Unbound | Mirror | 0.29 | 0.001 |
| PBP3 | Unbound | Native | 0.16 | 0.016 |
| PBP3 | Unbound | Mirror | 0.38 | 0.012 |
| IleRS | Unbound | Native | 0.29 | 0.028 |
| IleRS | Unbound | Mirror | 0.35 | 0.001 |
| MurA | Fosfomycin | Native | 0.15 | 0.012 |
| MurA | Fosfomycin | Mirror | 0.45 | 0.167 |
| DHPS | Sulfamethoxazole | Native | 0.19 | 0.003 |
| DHPS | Sulfamethoxazole | Mirror | 0.28 | 0.005 |
| PBP3 | Amoxicillin | Native | 0.27 | 0.02 |
| PBP3 | Amoxicillin | Mirror | 0.31 | 0.022 |
| PBP3 | Cephalexin | Native | 0.26 | 0.004 |
| PBP3 | Cephalexin | Mirror | 0.38 | 0.024 |
| IleRS | Mupirocin | Native | 0.29 | 0.08 |
| IleRS | Mupirocin | Mirror | 0.31 | 0.034 |
| PBP3 | Mirror-Amoxicillin | Native | 0.34 | 0.06 |
| PBP3 | Mirror-Amoxicillin | Mirror | 0.23 | 0.04 |

**SI Table 2.** Radius of Gyration.

| Protein | Enantiomer | RoG (nm) | StDev |
| --- | --- | --- | --- |
| MurA | Native | 2.23 | 0.004 |
| MurA | Mirror | 2.31 | 0.019 |
| DHPS | Native | 2.68 | 0.012 |
| DHPS | Mirror | 2.65 | 0.008 |
| PBP3 | Native | 1.98 | 0.012 |
| PBP3 | Mirror | 1.97 | 0.015 |
| IleRS | Native | 3.46 | 0.002 |
| IleRS | Mirror | 3.46 | 0.003 |

**SI Table 3.**
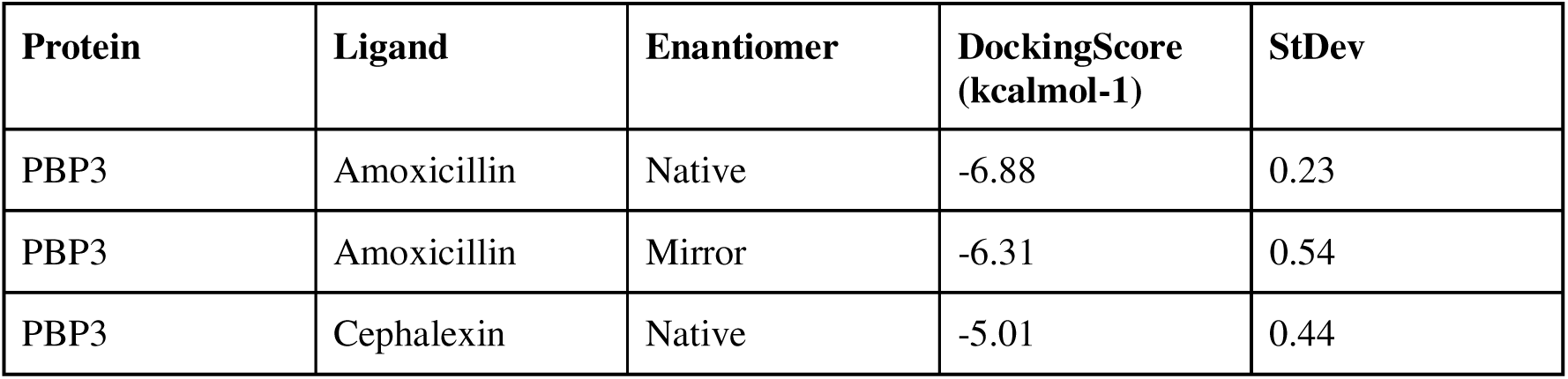
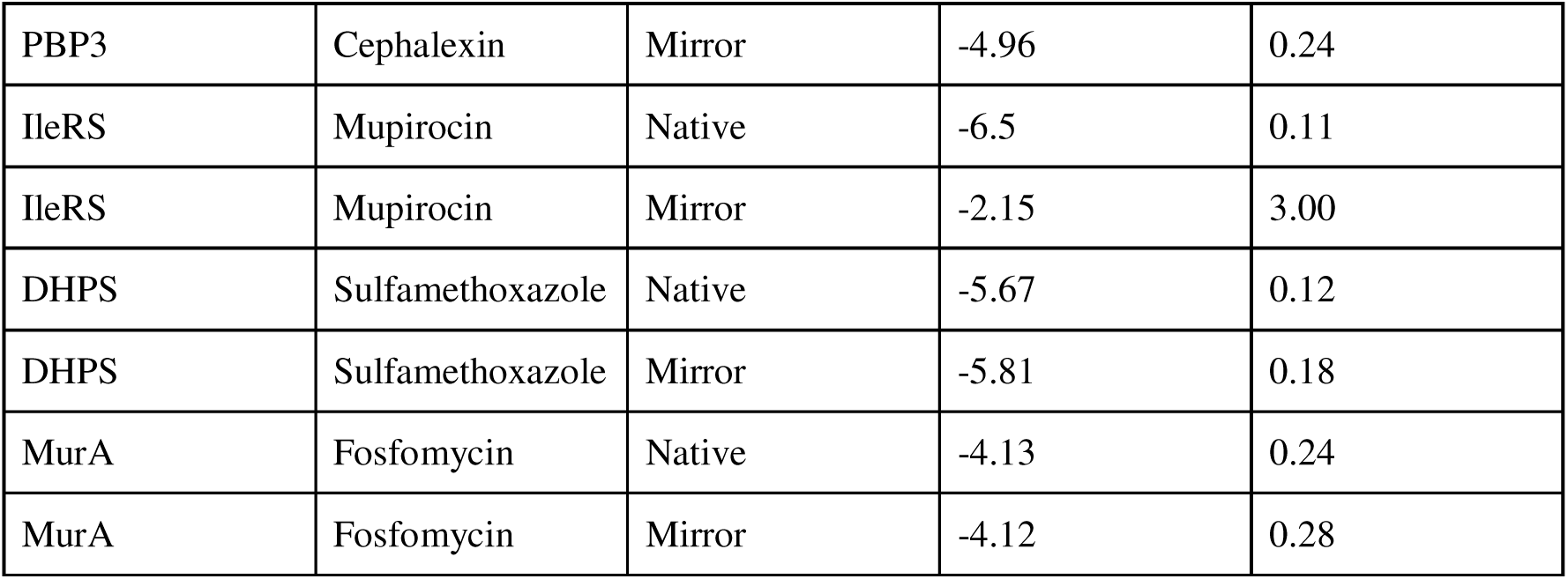
Docking Scores.

**SI Table 4.** Key Residue Distance.

| Protein | Ligand | Enantiomer | KeyResidue | Distance (nm) | StDev |
| --- | --- | --- | --- | --- | --- |
| MurA | Fosfomycin | Native | CYS_115 | 1.11 | 0.062 |
| MurA | Fosfomycin | Mirror | CYS_115 | 2.06 | 0.27 |
| DHPS | Sulfamethoxazole | Native | THR_51 | 1.4 | 0.021 |
| DHPS | Sulfamethoxazole | Mirror | THR_51 | 1.95 | 0.05 |
| PBP3 | Amoxicillin | Native | SER_320 | 0.99 | 0.3 |
| PBP3 | Amoxicillin | Mirror | SER_320 | 5.54 | 0.28 |
| PBP3 | Cephalexin | Native | SER_320 | 1.31 | 0.53 |
| PBP3 | Cephalexin | Mirror | SER_320 | 4.67 | 1.38 |
| IleRS | Mupirocin | Mirror | LEU_583 | 1.28 | 0.19 |
| IleRS | Mupirocin | Native | LEU_583 | 1.71 | 0.012 |
| PBP3 | Mirror-Amoxicillin | Native | SER_320 | 2.05 | 1.25 |
| PBP3 | Mirror-Amoxicillin | Mirror | SER_320 | 1.72 | 1.9 |

## References

1. Seeberger, P. H. Essentials of Glycobiology: Chapter 2: Monosaccharide Diversity. (Cold Spring Harbor laboratory press, Cold Spring Harbor, New York, 2022).

2. Winogradoff, D. et al. Chiral Systems Made from DNA. Adv. Sci. 8, 2003113 (2021).

3. Berg, J. M., Tymoczko, J. L. & Stryer, L. Biochemistry. (W. H. Freeman and company, New York, 2007).

4. Franklin, T. J. & Snow, G. A. Vulnerable shields — the cell walls of bacteria and fungi. in Biochemistry and Molecular Biology of Antimicrobial Drug Action 15–41 (Springer Netherlands, Dordrecht, 1998). doi:10.1007/978-94-010-9127-5_2.

5. Adamala, K. et al. Technical Report on Mirror Bacteria: Feasibility and Risks. 10.25740/CV716PJ4036 (2024) doi:10.25740/CV716PJ4036.

6. The Spirit of Asilomar and the Future of Biotechnology summit. 4.4 Risks from Mirror Life. https://hdl.handle.net/1911/118559 (2025) doi:10.25611/X89A-D947.

7. Paris Conference on Risks from Mirror Life. Paris Conference on Risks from Mirror Life: Meeting Report. https://zenodo.org/records/17167205 (2025) doi:10.5281/zenodo.17167205.

8. Zentrale Kommission für die Biologische Sicherheit. Spiegelbakterien. https://zkbs-online.de/synthetische-biologie/spiegelbakterien (2025).

9. International Bioethics Committee. Draft Report of the International Bioethics Committee (IBC) on the Ethical Issues Related to the Research, Development and Application of Synthetic Biology. https://unesdoc.unesco.org/ark:/48223/pf0000395064 (2025).

10. Kleinman, A., Torres, J. & Wang, B. Evaluation of antibiotic and peptide vaccine strategies for mirror bacterial infections. 2025.10.10.681688 Preprint at 10.1101/2025.10.10.681688 (2025).

11. Pedroni, L., Galaverna, G., Dall’Asta, C. & Dellafiora, L. Computational Investigations on Structural Aspects of Mirror Life. Preprint at 10.5281/ZENODO.14672236 (2025).

12. Pedroni, L., Dall’Asta, C., Galaverna, G. & Dellafiora, L. Computational Perspectives on Amoxicillin and Staphylococcus Aureus in Mirror Life. Glob. Chall. Hoboken NJ 9, e00051 (2025).

13. Burley, S. K. et al. Protein Data Bank (PDB): The Single Global Macromolecular Structure Archive. in Protein Crystallography (eds Wlodawer, A., Dauter, Z. & Jaskolski, M.) vol. 1607 627–641 (Springer New York, New York, NY, 2017).

14. Schonbrunn, E. E. cloacae MurA dead-end complex with UNAG and fosfomycin: 3lth. 10.2210/pdb3lth/pdb (2010).

15. Kostrewa, D., Oefner, C. & D’Arcy, A. DIHYDROPTEROATE SYNTHETASE (APO FORM) FROM STAPHYLOCOCCUS AUREUS: 1ad1. 10.2210/pdb1ad1/pdb (1998).

16. Bellini, D., Koekemoer, L., Newman, H. & Dowson, C. G. Crystal structure of TP domain from Chlamydia trachomatis Penicillin-Binding Protein 3 in complex with amoxicillin: 6i1f. 10.2210/pdb6i1f/pdb (2019).

17. Nakama, T., Nureki, O., Yokoyama, S., & RIKEN Structural Genomics/Proteomics Initiative (RSGI). Isoleucyl-tRNA synthetase Complexed with mupirocin: 1jzs. 10.2210/pdb1jzs/pdb (2001).

18. Brosnan, J. T. & Brosnan, M. E. The sulfur-containing amino acids: an overview. J. Nutr. 136, 1636S–1640S (2006).

19. Van Der Spoel, D. et al. GROMACS: Fast, flexible, and free. J. Comput. Chem. 26, 1701–1718 (2005).

20. Brooks, B. R. et al. CHARMM: The biomolecular simulation program. J. Comput. Chem. 30, 1545–1614 (2009).

21. Kim, S. et al. PubChem 2025 update. Nucleic Acids Res. 53, D1516–D1525 (2025).

22. Santos-Martins, D. et al. Meeko: Molecule Parametrization and Software Interoperability for Docking and Beyond. J. Chem. Inf. Model. 65, 13045–13050 (2025).

23. Yun, M.-K. et al. Catalysis and Sulfa Drug Resistance in Dihydropteroate Synthase. Science 335, 1110–1114 (2012).

24. Zhang, Y. & Skolnick, J. TM-align: a protein structure alignment algorithm based on the TM-score. Nucleic Acids Res. 33, 2302–2309 (2005).

25. Eberhardt, J., Santos-Martins, D., Tillack, A. F. & Forli, S. AutoDock Vina 1.2.0: New Docking Methods, Expanded Force Field, and Python Bindings. J. Chem. Inf. Model. 61, 3891–3898 (2021).

26. Bhati, S. K. et al. In silico screening and molecular dynamics analysis of natural DHPS enzyme inhibitors targeting Acinetobacter baumannii. Sci. Rep. 15, 7723 (2025).

27. Vanommeslaeghe, K. et al. CHARMM general force field: A force field for drug like molecules compatible with the CHARMM all atom additive biological force fields. J. Comput. Chem. 31, 671–690 (2010).

28. Wieczór, M., Czub, J. & Orozco, M. Gromologist: A GROMACS-oriented utility library for structure and topology manipulation. SoftwareX 30, 102118 (2025).

29. Adasme, M. F. et al. PLIP 2021: expanding the scope of the protein–ligand interaction profiler to DNA and RNA. Nucleic Acids Res. 49, W530–W534 (2021).

30. Laskowski, R. A. & Swindells, M. B. LigPlot+: Multiple Ligand–Protein Interaction Diagrams for Drug Discovery. J. Chem. Inf. Model. 51, 2778–2786 (2011).

31. Humphrey, W., Dalke, A. & Schulten, K. VMD: Visual molecular dynamics. J. Mol. Graph. 14, 33–38 (1996).

32. Ihaka, R. & Gentleman, R. R: A Language for Data Analysis and Graphics. J. Comput. Graph. Stat. 5, 299–314 (1996).

33. Wickham, H. ggplot2. WIREs Comput. Stat. 3, 180–185 (2011).

34. Osorio, D., Rondón-Villarreal, P. & Torres, R. Peptides: A Package for Data Mining of Antimicrobial Peptides. R J. 7, 4 (2015).

35. Bion, R. ricardo-bion/ggradar. (2025).

36. Grant, B. J., Rodrigues, A. P. C., ElSawy, K. M., McCammon, J. A. & Caves, L. S. D. Bio3d: an R package for the comparative analysis of protein structures. Bioinformatics 22, 2695–2696 (2006).

37. Minato Nakazawa, fmsb: Functions for Medical Statistics Book with some Demographic Data. 0.7.6 10.32614/CRAN.package.fmsb (2010).

38. Daina, A., Michielin, O. & Zoete, V. SwissADME: a free web tool to evaluate pharmacokinetics, drug-likeness and medicinal chemistry friendliness of small molecules. Sci. Rep. 7, 42717 (2017).

39. Cheng, T. et al. Computation of Octanol−Water Partition Coefficients by Guiding an Additive Model with Knowledge. J. Chem. Inf. Model. 47, 2140–2148 (2007).

40. Prasanna, S. & Doerksen, R. J. Topological polar surface area: a useful descriptor in 2D-QSAR. Curr. Med. Chem. 16, 21–41 (2009).

41. Delaney, J. S. ESOL: Estimating Aqueous Solubility Directly from Molecular Structure. J. Chem. Inf. Comput. Sci. 44, 1000–1005 (2004).

42. SanthaKumari, S. L. G. N. & Vincent, S. G. Identification and molecular characterization of drug targets of methicillin resistant Staphylococcus aureus. J. Appl. Nat. Sci. 14, 1152–1157 (2022).

43. Batista, F. A. et al. Oligomeric protein interference validates druggability of aspartate interconversion in Plasmodium falciparum. MicrobiologyOpen 8, e00779 (2019).

44. Wimley, W. C. & White, S. H. Experimentally determined hydrophobicity scale for proteins at membrane interfaces. Nat. Struct. Biol. 3, 842–848 (1996).

45. Najafi, S., Lobo, S., Shell, M. S. & Shea, J.-E. Context Dependency of Hydrophobicity in Intrinsically Disordered Proteins: Insights from a New Dewetting Free Energy-Based Hydrophobicity Scale. J. Phys. Chem. B 129, 1904–1915 (2025).

46. Boman, H. G. Antibacterial peptides: basic facts and emerging concepts. J. Intern. Med. 254, 197–215 (2003).

47. Han, H. et al. The Fungal Product Terreic Acid Is a Covalent Inhibitor of the Bacterial Cell Wall Biosynthetic Enzyme UDP-N-Acetylglucosamine 1-Carboxyvinyltransferase (MurA),. Biochemistry 49, 4276–4282 (2010).

48. Baum, E. Z. et al. Identification and Characterization of New Inhibitors of the Escherichia coli MurA Enzyme. Antimicrob. Agents Chemother. 45, 3182–3188 (2001).

49. Mezzatesta, M. L., Gona, F. & Stefani, S. Enterobacter cloacae Complex: Clinical Impact and Emerging Antibiotic Resistance. Future Microbiol. 7, 887–902 (2012).

50. Musil, I., Jensen, V., Schilling, J., Ashdown, B. & Kent, T. Enterobacter cloacae infection of an expanded polytetrafluoroethylene femoral-popliteal bypass graft: a case report. J. Med. Case Reports 4, 131 (2010).

51. Frutos-Grilo, E., Kreling, V., Hensel, A. & Campoy, S. Host-pathogen interaction: Enterobacter cloacae exerts different adhesion and invasion capacities against different host cell types. PLOS ONE 18, e0289334 (2023).

52. Sanders, W. E. & Sanders, C. C. Enterobacter spp.: pathogens poised to flourish at the turn of the century. Clin. Microbiol. Rev. 10, 220–241 (1997).

53. Vander Meersche, Y., Cretin, G., Gheeraert, A., Gelly, J.-C. & Galochkina, T. ATLAS: protein flexibility description from atomistic molecular dynamics simulations. Nucleic Acids Res. 52, D384–D392 (2024).

54. Lemkul, J. A. Introductory Tutorials for Simulating Protein Dynamics with GROMACS. J. Phys. Chem. B 128, 9418–9435 (2024).

55. Kahan, F. M., Kahan, J. S., Cassidy, P. J. & Kropp, H. The Mechanism of Action of Fosfomycin (phosphonomycin). Ann. N. Y. Acad. Sci. 235, 364–386 (1974).

56. Yoon, H.-J. et al. Crystal structure of UDP-N-acetylglucosamine enolpyruvyl transferase from Haemophilus influenzae in complex with UDP-N-acetylglucosamine and fosfomycin. Proteins Struct. Funct. Bioinforma. 71, 1032–1037 (2008).

57. Wanke, C. & Amrhein, N. Evidence that the reaction of the UDP-N-acetylglucosamine 1-carboxyvinyltransferase proceeds through the O-phosphothioketal of pyruvic acid bound to Cys115 of the enzyme. Eur. J. Biochem. 218, 861–870 (1993).

58. Evidence That the Fosfomycin Target Cys115 in UDP-N-acetylglucosamine Enolpyruvyl Transferase (MurA) Is Essential for Product Release − ScienceDirect. https://www.sciencedirect.com/science/article/pii/S0021925820762460.

59. Bensen, D. C., Rodriguez, S., Nix, J., Cunningham, M. L. & Tari, L. W. Structure of MurA (UDP-N-acetylglucosamine enolpyruvyl transferase) from Vibrio fischeri in complex with substrate UDP-N-acetylglucosamine and the drug fosfomycin. Acta Crystallograph. Sect. F Struct. Biol. Cryst. Commun. 68, 382–385 (2012).

60. Lima, A. H., Silva, J. R. A., Alves, C. N. & Lameira, J. QM/MM Study of the Fosfomycin Resistance Mechanism Involving FosB Enzyme. ACS Omega 6, 12507–12512 (2021).

61. Silver, L. L. Fosfomycin: Mechanism and Resistance. Cold Spring Harb. Perspect. Med. 7, a025262 (2017).

62. Hampele, I. C. et al. Structure and function of the dihydropteroate synthase from *staphylococcus aureus*1. J. Mol. Biol. 268, 21–30 (1997).

63. Sanyal, D. et al. Mapping dihydropteroate synthase evolvability through identification of a novel evolutionarily critical substructure. Int. J. Biol. Macromol. 311, 143325 (2025).

64. Arputharaj, D. S., Rajasekaran, M. & Nidhin, P. V. Sulfamethoxazole: Molecular docking and crystal structure prediction. Results Chem. 5, 100716 (2023).

65. Hammoudeh, D. I. et al. Identification and Characterization of an Allosteric Inhibitory Site on Dihydropteroate Synthase. ACS Chem. Biol. 9, 1294–1302 (2014).

66. Griffith, E. C. et al. The Structural and Functional Basis for Recurring Sulfa Drug Resistance Mutations in Staphylococcus aureus Dihydropteroate Synthase. Front. Microbiol. 9, (2018).

67. Griffith, E. C. et al. The Structural and Functional Basis for Recurring Sulfa Drug Resistance Mutations in Staphylococcus aureus Dihydropteroate Synthase. Front. Microbiol. 9, 1369 (2018).

68. Ploetz, E. A., Kariyawasam, N. L. & Smith, P. E. Probing Contributions to the Volume and Isothermal Compressibility of Globular Proteins Using a Fluctuation Solution Theory Analysis of MD Simulations. J. Phys. Chem. B 129, 8281–8298 (2025).

69. Ghuysen, J.-M. & Goffin, C. Lack of Cell Wall Peptidoglycan versus Penicillin Sensitivity: New Insights into the Chlamydial Anomaly. Antimicrob. Agents Chemother. 43, 2339–2344 (1999).

70. Skilton, R. J. et al. Penicillin Induced Persistence in Chlamydia trachomatis: High Quality Time Lapse Video Analysis of the Developmental Cycle. PLOS ONE 4, e7723 (2009).

71. Kocaoglu, O., Tsui, H.-C. T., Winkler, M. E. & Carlson, E. E. Profiling of β-Lactam Selectivity for Penicillin-Binding Proteins in Streptococcus pneumoniae D39. Antimicrob. Agents Chemother. 59, 3548–3555 (2015).

72. Houba-Hérin, N., Hara, H., Inouye, M. & Hirota, Y. Binding of penicillin to thiol-penicillin-binding protein 3 of Escherichia coli: identification of its active site. Mol. Gen. Genet. MGG 201, 499–504 (1985).

73. Sauvage, E., Kerff, F., Terrak, M., Ayala, J. A. & Charlier, P. The penicillin-binding proteins: structure and role in peptidoglycan biosynthesis. FEMS Microbiol. Rev. 32, 234–258 (2008).

74. Kocaoglu, O. & Carlson, E. E. Profiling of β-Lactam Selectivity for Penicillin-Binding Proteins in Escherichia coli Strain DC2. Antimicrob. Agents Chemother. 59, 2785–2790 (2015).

75. Nakama, T., Nureki, O. & Yokoyama, S. Structural Basis for the Recognition of Isoleucyl-Adenylate and an Antibiotic, Mupirocin, by Isoleucyl-tRNA Synthetase*. J. Biol. Chem. 276, 47387–47393 (2001).

76. Chung, S., Kim, S., Ryu, S. H., Hwang, K. Y. & Cho, Y. Structural Basis for the Antibiotic Resistance of Eukaryotic Isoleucyl-tRNA Synthetase. Mol. Cells 43, 350–359 (2020).

77. Zanki, V., Bozic, B., Mocibob, M., Ban, N. & Gruic-Sovulj, I. A pair of isoleucyl-tRNA synthetases in Bacilli fulfills complementary roles to keep fast translation and provide antibiotic resistance. Protein Sci. Publ. Protein Soc. 31, e4418 (2022).

78. Murthy, S. et al. Biological Characterization of Mupirocin–KGF Hydrogel and Its Regenerative Potential in Human Fibroblast-Mediated Wound Healing. Molecules 30, 4523 (2025).

79. Závodszky, P., Kardos, J., Svingor, Á. & Petsko, G. A. Adjustment of conformational flexibility is a key event in the thermal adaptation of proteins. Proc. Natl. Acad. Sci. 95, 7406–7411 (1998).

80. The Survival Mechanisms of Thermophiles at High Temperatures: An Angle of Omics | Physiology | American Physiological Society. Physiology https://journals.physiology.org/doi/10.1152/physiol.00066.2013.

81. Najmanovich, R., Kuttner, J., Sobolev, V. & Edelman, M. Side-chain flexibility in proteins upon ligand binding. Proteins 39, 261–268 (2000).

82. Carlson, H. A. Protein flexibility and drug design: how to hit a moving target. Curr. Opin. Chem. Biol. 6, 447–452 (2002).

83. Ligand flexibility and binding pocket malleability cooperate to allow selective PXR activation by analogs of a promiscuous nuclear receptor ligand −ScienceDirect. https://www.sciencedirect.com/science/article/pii/S0969212623002976.

84. Perspectives on Ligand/Protein Binding Kinetics Simulations: Force Fields, Machine Learning, Sampling, and User-Friendliness | Journal of Chemical Theory and Computation. https://pubs.acs.org/doi/10.1021/acs.jctc.3c00641.

85. Grewal, S., Deswal, G., Grewal, A. S. & Guarve, K. Molecular dynamics simulations: Insights into protein and protein ligand interactions. Adv. Pharmacol. 103, 139–162 (2025).

86. Singh, J., Petter, R. C., Baillie, T. A. & Whitty, A. The resurgence of covalent drugs. Nat. Rev. Drug Discov. 10, 307–317 (2011).

87. De Vita, E. 10 Years Into the Resurgence of Covalent Drugs. Future Med. Chem. 13, 193–210 (2021).

88. Mora-Ochomogo, M. & Lohans, C. T. β-Lactam antibiotic targets and resistance mechanisms: from covalent inhibitors to substrates. RSC Med. Chem. 12, 1623–1639 (2021).

89. Carro, L. Recent Progress in the Development of Small-Molecule FtsZ Inhibitors as Chemical Tools for the Development of Novel Antibiotics. Antibiotics 8, 217 (2019).

90. Das, S., Poudel, R., Dutta, K. & Konai, M. M. A review on small molecular mimics of antimicrobial peptides with an emphasis on the structure–activity relationship perspective. RSC Med. Chem. 16, 3982–4002 (2025).

91. Kabsch, W. & Sander, C. Dictionary of protein secondary structure: Pattern recognition of hydrogen-bonded and geometrical features. Biopolymers 22, 2577–2637 (1983).

